# The molecular chronology of mammary epithelial cell fate switching

**DOI:** 10.1101/2024.10.08.617155

**Authors:** Queralt Vallmajo-Martin, Zhibo Ma, Sumana Srinivasan, Divya Murali, Christopher Dravis, Kavitha Mukund, Shankar Subramaniam, Geoffrey M. Wahl, Nikki K. Lytle

## Abstract

The adult mammary gland is maintained by lineage-restricted progenitor cells through pregnancy, lactation, involution, and menopause. Injury resolution, transplantation-associated mammary gland reconstitution, and tumorigenesis are unique exceptions, wherein mammary basal cells gain the ability to reprogram to a luminal state. Here, we leverage newly developed cell-identity reporter mouse strains, and time-resolved single-cell epigenetic and transcriptomic analyses to decipher the molecular programs underlying basal-to-luminal fate switching *in vivo*. We demonstrate that basal cells rapidly reprogram toward plastic cycling intermediates that appear to hijack molecular programs we find in bipotent fetal mammary stem cells and puberty-associatiated cap cells. Loss of basal-cell specifiers early in dedifferentiation coincides with activation of Notch and BMP, among others. Pharmacologic blockade of each pathway disrupts basal-to-luminal transdifferentiation. Our studies provide a comprehensive map and resource for understanding the coordinated molecular changes enabling terminally differentiated epithelial cells to transition between cell lineages and highlights the stunning rapidity by which epigenetic reprogramming can occur in response to disruption of tissue structure.

## Introduction

Mammary glands, the defining feature and source for the name of the class “mammalia”, provide an excellent system to decipher mechanisms of cellular dynamics and plasticity. During murine development, fetal cells originating from the overlying epidermis generate a “mammary bulb” by embryonic day 12.5 (E12.5)^1, 2^. After a “resting period” fetal mammary progenitors proliferate extensively to generate primitive ducts composed of epithelial cells, the majority of which co-express transcripts found in adult luminal and basal cells thereby exhibiting a “hybrid cell state”^3–6^.

Shortly after birth, rapid specification occurs into two broad luminal cell types comprising hormone sensing (i.e., “HS”; formerly mature luminal/ML) and alveolar luminal secretory cells (i.e., “LS”; formerly luminal progenitor/LP), and an external sheath of muscle-like basal/myoepithelial cells. Ductal elongation, which continues until puberty, is achieved by luminal cells and basal-like cap cells, which terminate the outer layer of terminal end-buds (TEBs) and enable ducts to extend through the fat pad^7, 8^. Basal-layer cap cells of the TEBs have been proposed to be capable of generating underlying luminal cells^7–9^, but murine TEBs and cap cells disappear after puberty, resulting in blunt-ended homeostatic adult ducts^7–9^. This contrasts with mammary ducts in humans and rats, which are terminated by multiple complex lobular structures referred to as terminal duct lobular units (TDLUs)^10^. In adult female mice, mammary glands undergo exuberant proliferation in response to pregnancy hormones to generate alveoli that produce milk during lactation. After weaning, involution, a process involving induction of apoptosis on a mass scale followed by remodeling of the epithelium, restores the mammary gland to a state resembling that prior to pregnancy^11, 12^.

Early studies suggested that mammary gland homeostasis may be achieved by stem cells originating embryonically that persist into adulthood. This was supported by studies demonstrating that fetal mammary cells and a subset of basal adult mammary epithelial cells generate full, functional mammary glands when transplanted into cleared fat pads^13, 14^, a cardinal characteristic of “stem cells”^9, 15–17^. However, elegant lineage tracing studies, which leveraged indelible marks targeted toward specific cell populations in situ, resulted in discrepent findings^5, 18–27^. They confirmed the existence of bipotent fetal mammary stem cells (fMaSCs)^5, 6, 19, 20, 24^ while also demonstrating that adult basal, LS, and HS cell populations are maintained by unipotent, lineage-restricted progenitors during puberty, pregnancy, lactation, and involution^18, 21, 25–27^. While two lineage tracing studies suggest that a small population of basal cells may be bipotent mammary stem cells (MaSCs) in homeostatic glands^9, 28^, saturation lineage tracing and mathematical modeling found no evidence of such cells^5, 18–21^. Resolving the apparent difference between cell potential evaluated by transplantation and cell fate established by lineage tracing remains a major unresolved question.

Here, we test the hypothesis that cell state reprogramming of basal cells to a multipotent state in response to tissue structure disruption underlies their ability to regenerate functional mammary glands following transplantation. This hypothesis is consistent with transplantation studies showing that disrupting tissue homeostasis can lead to cell fate reprogramming in lung^29^ and skin^30–32^, as can selective diptheria toxin induced ablation of mammary luminal cells^33^.

Further, chemotherapy induced organ damage also enables mammary basal-to-luminal reprogramming in situ^34^. These studies reveal that a common strategy for wound repair in adult tissues may involve fully differentiated cells acquiring developmental plasticity as they are freed from inhibitory cell-cell and cell-matrix associations, and/or upon exposure to wound induced microenvironmental changes. However, the molecular mechanisms enabling acquired plasticity remain poorly understood.

In order to test this hypothesis and provide insights into the kinetics, molecular pathways, and cell state regulators involved, we developed unique mouse models that enable us to decipher the relevant transcriptomic and epigenetic changes at single cell resolution. We used CRISPR to introduce green and red nuclear-localized fluorescent proteins immediately ahead of the stop codon and the 3’ UTRs of the keratin 14, 8, 18, and 19 genes. As the fluorescent proteins are controlled by the endogenous regulatory elements of these genes, this system enabled us to localize, isolate, and analyze cells in the basal (green), luminal (red), or transitioning (mixture of green and red) cell states. The results demonstrate that under conditions of transplantation, basal cells rapidly reprogram into a mixed lineage state with transcriptomic and epigenetic relatedness to both fMaSCs and cap cells that mediate ductal elongation in puberty. Thus, the mammary gland may be utilizing mechanisms of plasticity used in the cells from which it is developmentally derived to re-establish homeostasis.

## Results

### Development of cell-state fluorescent reporter mice enables tracking, isolation and molecular characterization of all mammary epithelial cell lineages

Cell surface markers compatible with fluorescence-activated cell sorting (FACS) have historically been used to isolate mammary epithelial cell lineages for molecular and functional studies. These markers, including EpCAM^35^, Integrin alpha-6 (CD49f)^36^, CD24^16, 37^, Integrin beta-1 (CD29)^16^, Integrin beta-3 (CD61)^38^, Integrin alpha-2 (CD49b)^39^, cKit^40^, and Integrin alpha-3, correlate with fully differentiated cell lineages. We reasoned that identifying unstable cell states that may arise during mammary regeneration, wound healing, oncogenesis, or other conditions would benefit from reporter mice using genes more tightly linked to cell state. We therefore created two basal cell state indicators based on Keratin 14 (K14) expression, K14-mClover and K14-tdTomato (K14-mCl and K14-tdT, respectively), and three luminal cell state indicators, Keratin8-tdTomato, Keratin18-tdTomato, and Keratin19-mClover (K8-tdT, K18-tdT, and K19-mCl, respectively) (**Figure 1A, Figure S1A**). In all mouse models, we used CRISPR to insert a self-cleaving P2A sequence followed by a fluorophore cassette adjacent to the 3’ termination of the endogenous keratin locus. Flow cytometry analysis of each reporter revealed robust overlap with commonly used cell surface staining used to identify mammary epithelial cell populations; e.g. EpCAM^lo^/CD49f^Hi^ for basal cells and EpCAM^Hi^;CD49f^Lo^ for luminal cells (**Figure S1B-G**). Analysis of K14 fluorophore expression revealed that the fluorescent knock-in alleles (K14-tdT and K14-mCl) reported basal lineage more accurately than a previously reported K14-RFP transgenic mouse generated using the K14 promoter driving fluorophore expression^41^ (**Figure S1H-I**). This is consistent with enhancer elements in the native locus being critical for faithful gene expression. Further, reporter expression in developing embryonic mammary buds was visible with a fluorescent dissecting microscope in intact embryos as early as embryonic day 12.5 (E12.5) and in skin flanks of later stage embryos (**Figure 1B**). This proved useful for isolating and analyzing mammary epithelium during development^42^.

**Figure 1.**
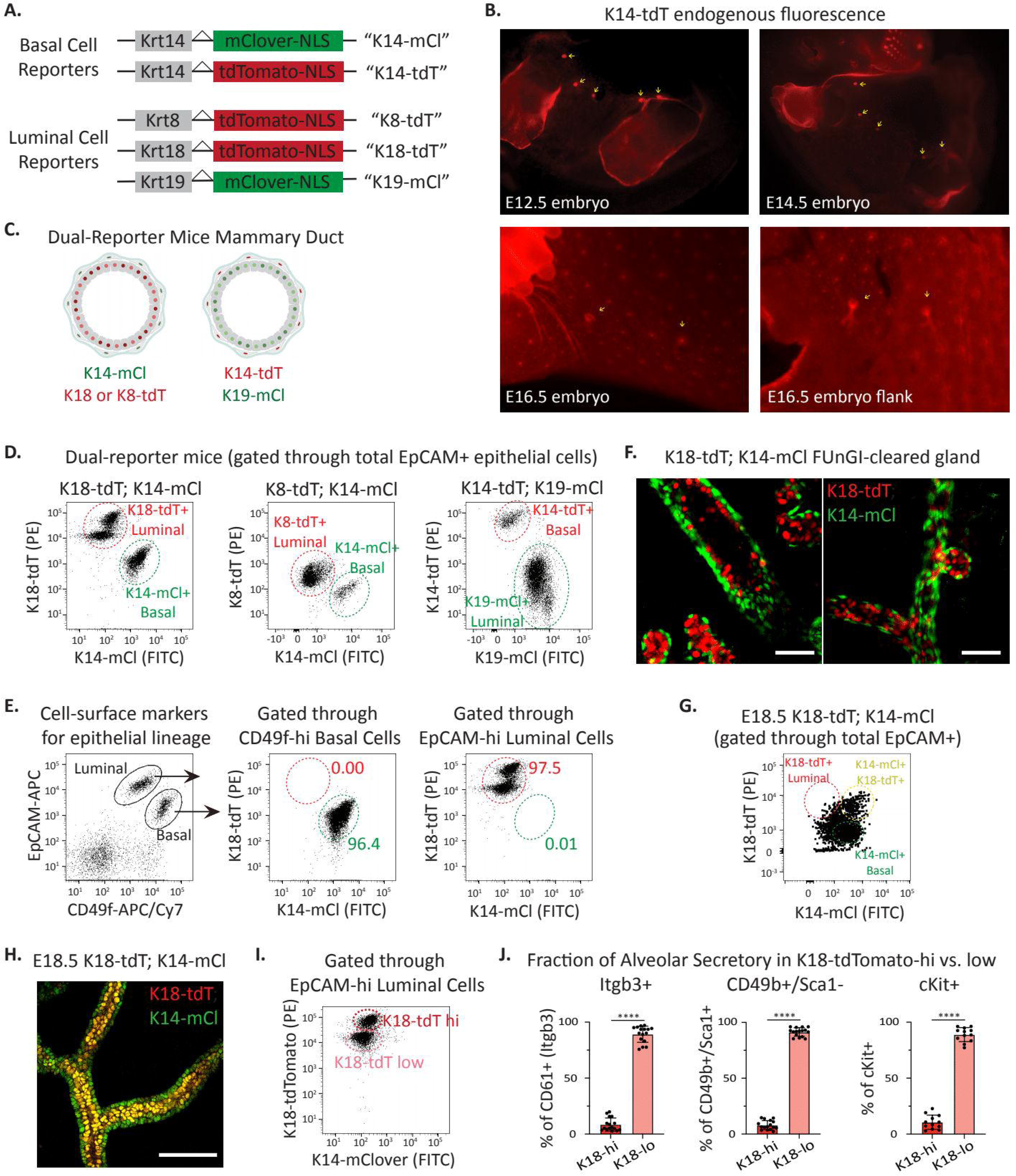
Development of cell-state fluorescent reporter mice enables tracking, isolation and molecular characterization of all mammary epithelial cell lineages. A. Design of mammary epithelial cell state indicator mice. Basal cell reporters: Keratin14 (Krt14)–mClover (K14-mCl) and Krt14-tdTomato (K14-tdT). Luminal cell reporters: K8-tdT, K18-tdT, and K19-mCl. Self-cleaving 2A sequences separate each keratin from the fluorophore. B. K14-tdT endogenous red fluorescence of female mouse embryos at embryonic day 12.5 (E12.5), E14.5, and E16.5 whole embryos, and E16.5 skin flanks. Yellow arrows indicate developing mammary buds. C. Schematic of dual-reporter mice mammary ducts. Basal and luminal cells are indicated by red or green fluorescence in K14-mCl; K18-tdT or K14-mCl; K8-tdT (left), or K14-tdT; K19-mCl (right). Created with BioRender.com. D. Flow cytometry analysis of adult dual-reporter mouse mammary epithelium. E. Flow cytometry analysis of adult K18-tdT; K14-mCl dual-reporter mammary gland. Left plot indicates gates identifying EpCAM^Hi^; CD49f^Lo^ luminal cells and CD49f^Hi^; EpCAM^Lo^ basal cells. K18-tdT and K14-mCl endogenous fluorescent expression is shown for basal cells (middle) and luminal cells (right). F. Representative images of endogenous fluorescence from FUnGI-cleared adult K18-tdT; K14-mCl dual-reporter mammary glands. Scale bars: 50 μm. G. Flow cytometry analysis of K18-tdT; K14-mCl dual-reporter mammary buds from E18.5 female embryos. Highlighted luminal cells (K18-tdT+), basal cells (K14-mCl+), and double-positive (++, yellow) gating within total epithelial cells (EpCAM^+^). H. Representative image of K18-tdT; K14-mCl dual-reporter mammary bud endogenous fluorescence from an E18.5 female embryo. Yellow cells indicate dual expression of K18-tdT and K14-mCl. Scale bar: 100 μm. I. Flow cytometry analysis of K18-tdT expression in EpCAM^Hi^; CD49f^Lo^ luminal cells. Two populations are indicated by K18-tdT hi and lo expression. J. Analysis of K18-tdT hi and lo expression within alveolar luminal secretory cells identified as EpCAM^Hi^; CD49f^Lo^ luminal and Itgb3+ (left), CD49b+/Sca1- (middle), or cKit+ (right). n = 12-16 per group. Data are represented as mean ± SD. Student’s t test; **** p ≤ 0.0001

We generated “dual-reporter mice” by crossing basal with luminal reporter mice expressing different fluorophores (**Figure 1C-D**). As anticipated from analysis of individual reporter mice, flow cytometry analysis of dual reporters indicated substantial overlap with expected cell-surface lineage markers (**Figure 1E**). To visualize reporter expression in adult mammary glands with single-cell resolution, we used FUnGI clearing^43^ of a K18-tdT; K14-mCl mammary gland from a nulliparous, adult mouse followed by 3D confocal imaging. This revealed lumen-facing K18-tdT expressing cells surrounded by basal K14-mCl+ cells (**Figure 1F**). K14-mCl+; K18-tdT+ co-expressing cells as well as basal and luminal lineage “leaning” cells were evident by flow cytometry analysis of reporter expression in E18.5 mammary rudiments (**Figure 1G**) and observed by confocal imaging by the co-expression of both endogenous fluorophores (**Figure 1H, S1J**). This is consistent with our epigenetic and gene expression evidence indicating that E18.5 fMaSCs, while largely undifferentiated, manifest some features of lineage specification^44^.

The luminal keratin reporters, particularly K18-tdT, displayed clear and distinct high and low keratin populations by flow cytometry (**Figure 1I, Figure S1F**). Comparison of the K18-tdT high and low populations against previously reported markers of LS cells, including Integrin beta-3 (Itgb3/CD61^+^), Integrin alpha-2 (CD49b^+^)/Sca-1-negative, and c-Kit^+^ markers revealed that K18-tdT-low luminal cells were significantly enriched for LS cell populations (**Figure 1J**; gating for each LS lineage shown in **Figure S1K**). These data demonstrate that our K18-reporter alone could be leveraged in lineage-tracing experiments in which LS or HS cells are tracked. For this reason, and the clear separation between red and green fluorophores, we primarily use K18-tdT; K14-mCl dual-reporter mice through most studies below.

We conducted single-cell RNA-sequencing (scRNA-seq) analysis on epithelial cells isolated by fluorescent-activated cell sorting (FACS) to assess the molecular similarity of epithelial populations from K18-tdT; K14-mCl knock-in reporters to those from wildtype mice. We used uniform manifold approximation and projection (UMAP) analysis to subcluster and visualize epithelial cell populations, and identified each cell cluster using previously published cell ID scores for basal, LS, and HS^44^. This revealed that reporter epithelial cell clusters significantly overlap with wildtype cell clusters, suggesting the transcriptional state of mammary epithelial cells is not altered by the knock-in reporter alleles (**Figure S1L)**. We conclude that the fluorescent knock-in keratin reporters accurately and effectively identify and characterize each different mammary epithelial cell type and preserve their transcriptomes.

### Basal and luminal keratin coexpressing cells do not represent adult mammary stem cells

Whether or not a mammary stem cell pool exists in the adult remains a significant question. The majority of *in vivo* lineage tracing experiments suggest that each cell type in the mammary gland is independently maintained by lineage-restricted progenitors through puberty, pregnancy, lactation, and involution^5, 18–21^. Notably, a few lineage tracing reports^9, 28^ and all transplant studies indicate that the basal cell compartment contains cells able to generate both basal and luminal cells. Interestingly, the dual reporter mice enabled us to isolate ∼0.2-1.6% of epithelial cells expressing both basal and luminal keratins in adults (**Figure 2A-B**). We reasoned that these cells may represent a rare resident bipotent stem cell population because they co-express basal and luminal cell type indicators, as do bipotent embryonic mammary stem cells^3, 6, 44^. To test if this population functionally exhibits stem-like features, we conducted *in vivo* mammary gland reconstitution functional assays using FACS-purified double-positive cells, basal cells, or luminal cells. These data revealed double-positive cells are functionally intermediate in their mammary reconstitution capacity relative to basal and luminal cells (**Figure 2C**). Flow cytometry analysis of cell surface marker expression on double-positive cells revealed the majority overlap with the luminal cell gate (EpCAM^Hi^/CD49f^Lo^) or fall between luminal and basal cell gates, and a smaller subset overlaps with the basal cell gate (CD49f^Hi^/EpCAM^Lo^) (**Figure 2D-E**), suggesting the double-positive population represents a mix of basal and luminal cells. This was further supported by *in vitro* 3D mammosphere forming assay. Pure basal cells give rise to mammospheres comprised of both basal and luminal cells while luminal cells give rise to cystic spheres largely comprised of luminal cells (**Figure 2F-H**). Double-positive cells form mammospheres with intermediate basal cell composition suggesting the structures may arise from either basal or luminal cells (**Figure 2F-H**).

**Figure 2.**
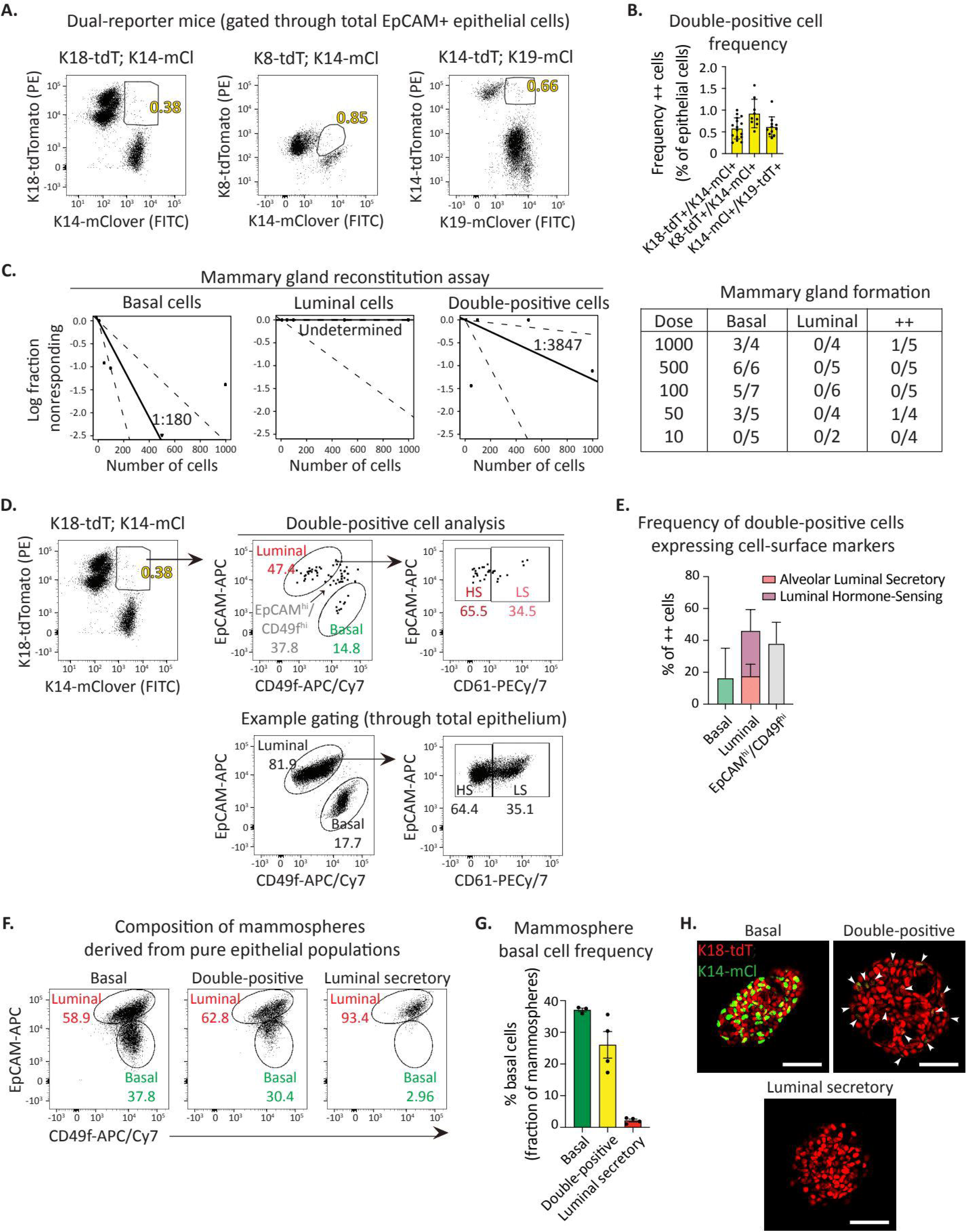
Basal and luminal keratin coexpressing cells do not represent adult mammary stem cells. A – B. Identification (A) and quantification (B) of adult mammary epithelial cells co-expressing reporters for basal (K14+) and luminal (K18+, K8+, or K19+) keratins by flow cytometry analysis. n = 9-16 per group. C. Mammary gland reconstitution assay from adult mammary epithelial cell populations. Limiting dilution frequency for mammary gland formation calculated for basal (left; K14-tdT+/K19-mCl-; 1:180), luminal (middle; K19-mCl+/K14-tdT-; undetermined), and double-positive (right; K14-tdT+/K19-mCl+; 1:3847). Table indicates cell doses tested in biological replicates. D. Flow cytometry analysis of K18-tdT+/K14-mCl+ double-positive (++) gating within basal (EpCAM^Lo^/CD49f^Hi^), LS (EpCAM^Hi^/CD49f^Lo^/CD61^+^), HS (EpCAM^Hi^/CD49f^Lo^/CD61^-^), or between basal and luminal gates (EpCAM^Hi^/CD49f^Hi^). E. Distribution of ++ cells in basal, LS, HS, and EpCAM^Hi^/CD49f^Hi^ gates. n = 15 K18-tdT; K14-mCl dual-reporter mice analyzed. F. Flow cytometry analysis of basal (EpCAM^Lo^ /CD49f^Hi^) and luminal (EpCAM^Hi^/CD49f^Lo^) cells comprising 3D mammospheres derived from pure basal (left; K14-mCl+/K18tdT-), double-positive (middle; K14-mCl+/K18-tdT+), or alveolar luminal secretory cells (right; K14-mCl-/K18-tdT+/CD61+). Mammospheres were dissociated and analyzed after two weeks in culture. G. Quantification of the frequency of basal cells comprising mammospheres derived from pure basal, double-positive, or alveolar luminal secretory cells. n = 3-4 biological replicates. H. Representative z-stack images of mammospheres derived from pure basal, double-positive, or alveolar luminal secretory cells two weeks in culture. Scale bars: 50 μm; z-stacks: 13 μm (basal) and 8 μm (double-positive and luminal secretory cells). White arrows indicate mClover+ cells in the mammospheres from double-positive cells. Data are represented as mean ± SD.

To determine if double positive cells molecularly resemble differentiated populations (e.g. basal, LS, or HS) or comprise a distinct cell cluster, we conducted scRNA-seq analysis on flow-sorted pure double positive cells as well as all other epithelial cells as reference populations. We employed UMAP analysis to visualize single-reporter positive epithelial cell populations, then overlayed the double-positive cells (**Figure S2A**). These data revealed that double-positive cells do not transcriptomically constitute a unique hybrid population, but rather resemble either basal, LS, or HS cells. Further, transcriptomically double-positive cells were not enriched for Tspan8^45^ or Procr^28^, two putative mammary stem cell markers (**Figure S2B**). We did note that double-positive cells are enriched for heat-shock related genes (**Figure S2C-D**), suggesting that co-expression of basal and luminal keratins may occur in response to stress-induced cell-state instability. Taken together, these data strongly suggest that cells co-expressing basal and luminal keratins are not tissue-resident stem cells.

### Basal cells rapidly acquire luminal cell characteristics *in vitro* and *in vivo*

Basal cells are uniquely able to reconstitute entire mammary glands upon transplantation *in vivo* (**Figure 2C**) and give rise to polarized mammospheres *in vitro* containing both an internal luminal cell layer and external basal cell layer (**Figure 2F-H**). Conversion of basal to luminal cells in mammosphere conditions has been reported to occur rapidly^46^, though the mechanisms remain unclear. The K18-tdT; K14-mCl dual-reporters provide a powerful system to ascertain the precise kinetics of basal-to-luminal cell state conversion and for deducing the molecular pathways involved. We isolated pure basal cells (K14-mCl+/K18-tdT-), plated them in 3D mammary mammosphere conditions and analyzed the resulting structures every 2 days. This analysis revealed that within days, basal cells began losing expression of K14-mCl to generate K18-tdT+ luminal cells (**Figure 3A-B**), highlighting the capability of this knock-in keratin reporter system to reveal cell state transitions under conditions that trigger cell state instability.

**Figure 3.**
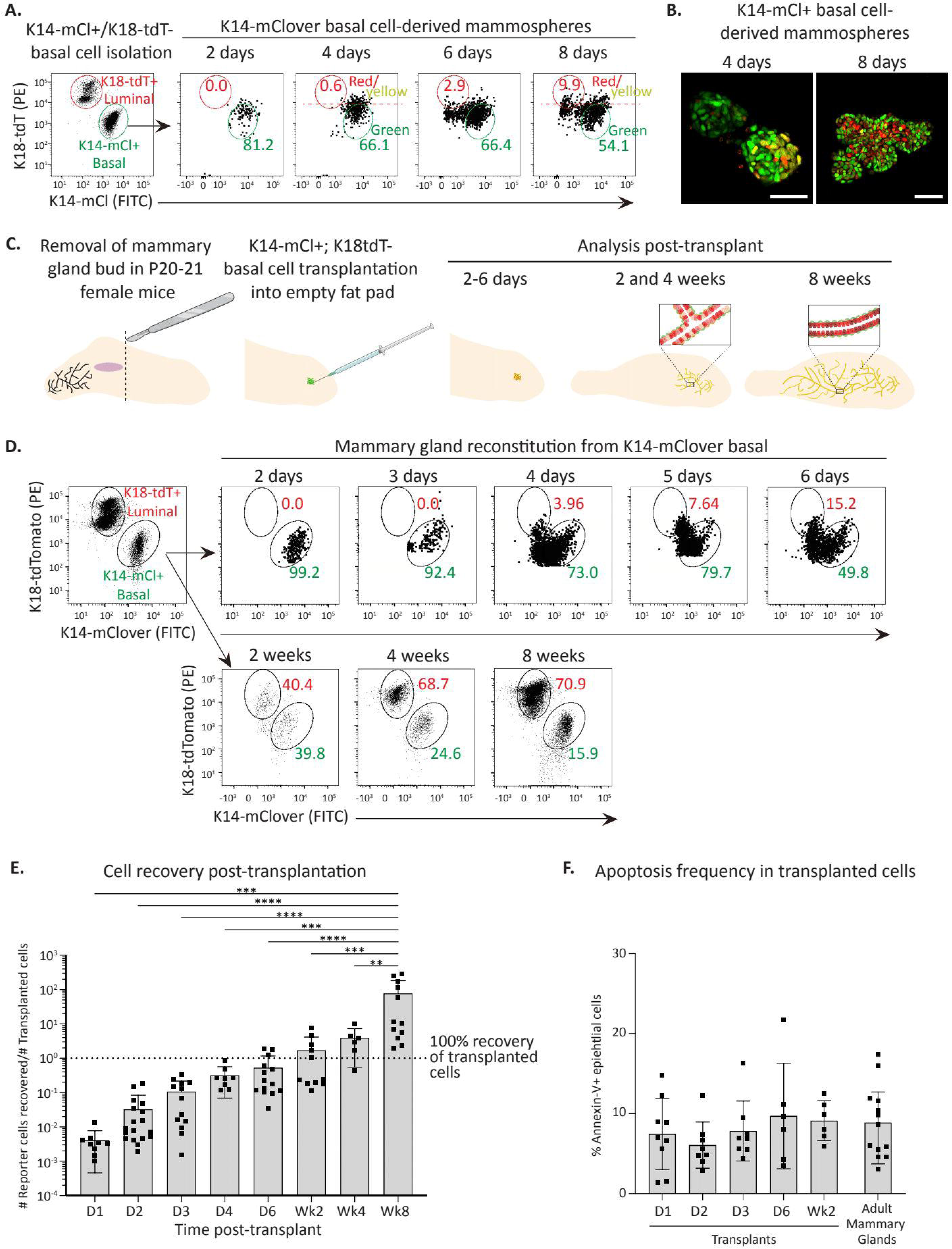
Basal cells rapidly acquire luminal cell characteristics *in vitro* and *in vivo*. A. Isolation of pure K14-mCl+; K18-tdT-basal cells (left plot) for mammosphere formation assay. Example flow plots of mammospheres at day 2, 4, 6, and 8 post-seeding. Cells above red dotted line will appear yellow or red by confocal imaging due to the emergence of tdTomato signal. B. Representative z-stack images of mammospheres derived from K14-mCl+ basal cells at 4 and 8 days after seeding. Nuclear expression of the K14-mCl (green), K18-tdT (red), or both (yellow). Scale bars: 50 μm; z-stacks: 30 μm (d4) and 11 μm (d8). C. Schematic of mammary gland reconstitution using pure basal cells from K14-mCl; K18-tdT adult female mice transplanted into de-epithelialized fat pads of postnatal day 20-21 female SCID mice. Cells were analyzed at days 2-6 and weeks 2, 4, and 8 post-transplant. Created with BioRender.com. D. Representative flow plots of reporter expression in transplant recipients at indicated time points. E. Total reporter+ epithelial cells recovered in transplant recipients over total transplanted cells at indicated time points. Dotted line indicates recovery of 100% of cells transplanted. n = 6-18 biological replicates per timepoint. F. Percentage of apoptotic cells expressing Annexin-V within total reporter+ epithelial cells in transplant recipients at indicated time points, or in murine adult mammary glands. n = 6-13 biological replicates per condition. Data are represented as mean ± SD. One-way ANOVA with Tukey’s multiple comparison test; ** p ≤ 0.01; *** p ≤ 0.001; **** p ≤ 0.0001.

We next determined if rapid basal-to-luminal fate switching occurs in vivo using mammary reconstitution assays in postnatal day 21 (P21) recipient SCID mice. Pure basal cells isolated from K14-mCl; K18-tdT adult dual-reporter mice were transplanted into de-epithelialized mammary fat pads, then analyzed at different timepoints post-transplant (**Figure 3C**). Strikingly, this kinetic analysis revealed that within 3-6 days, transplanted basal cells downregulated K14-mCl expression and began expressing K18-tdT (**Figure 3D**), though we noted significant biological variability between the exact day luminal reporter expression was observed (**Figure S3**). By 4 weeks post-transplant, distinct basal and luminal populations were observed (**Figure 3D, S3**). Mammary gland reconstitution is often measured 8 weeks post-transplant^13, 47^, and by this time we observed complete recapitulation of K18-tdT^Hi^ HS-enriched cells, K18-tdT^Lo^ LS-enriched cells, and K14-mCl+ basal cells (**Figure 3D, S3**).

We also tracked the cell numbers isolated at each time point and found a dramatic loss of cells following transplantation, cell number recovery by 6 days post-transplant, and increasing cell numbers until full reconstitution (**Figure 3E**). We next checked whether transplanted cells were undergoing apopotosis after transplantation. Annexin-V staining from recovered cells at early time points post-transplantation showed no significant difference compared to recovered cells from adult mice (**Figure 3F**). This indicates that initial transplanted cell loss is not due to basal cell apoptosis immediately after transplantation, but likely due to technical challenges during transplantation or cell re-isolation prior to analysis.

### Multiome analysis reveals molecular dynamics of basal-to-luminal cell state transitioning

The stunning rapidity with which basal cells begin expressing luminal keratins, and the fact that the basal cell epigenome most closely resembles that of fMaSCs^44^, suggests that basal cells are epigenetically poised to undergo luminal cell coversion under appropriate conditions. However, it has remained challenging to resolve the molecular mechanisms driving loss of basal identity and gain of luminal identity. Further, it remains unclear whether basal cells directly transdifferentiate to a luminal state or step through a less differentiated, perhaps fetal stem-like state. Our new reporter mice provide a powerful system to address these questions as it enables isolation and molecular characterization using single-cell epigenetic and transcriptomic analyses as well as functional validation of inferred pathways.

We performed four sets of analyses to develop an integrated cell state, epigenetic, and transcriptomic map representing the dynamics of basal to luminal conversion occurring in this system (**Figure 4A**). All studies used transplantion of K14-mCl+; K18-tdT-mammary basal epithelial cells isolated from adult dual-reporter mice (referred to as “**adult**”). We performed single cell analyses on this population to serve as a comparator for changes in differentiation state subsequent to transplantation. We first transplanted adult basal cells into de-epithelialized recipient mice, then re-isolated cells 3, 4, and 5 days after transplantion (“**D3-5**”). We combined all cells into a single pool for analysis to capture biologic and technical variability of cell transition observed in these early timepoints (**Figure S3**). We chose these time points as they encompass the earliest times at which we reproducibly observed basal cells losing basal keratin expression and gaining luminal keratin expression (**Figure 3D, S3**). In a second analysis, we transplanted basal cells and re-isolated them 6 days post-transplant (“**D6**”). This subset of cells consistently expressed luminal keratins, and therefore represents a time point for analyzing the earliest putative luminal cells (**Figure 3D, S3**). Finally, we analyzed reconstituted mammary glands that had developed entirely from transplanted basal cells at 8 weeks as by this time the transplanted cells had resolved into all lineages (“**Wk8**”). For all time points, we performed 10x Genomics Multiome to simultaneously profile chromatin accessibility (ATAC) and transcriptome (3’ RNA) from the same nucleus of each isolated cell (**Figure 4A**). After filtering for high quality nuclei, single nuclei, our final multiome analysis included: 5,006 nuclei from adult, 2,547 from D3-5, 10,166 from D6, and 3,556 from Wk8. While this is a powerful platform for gaining insights into the molecular framework underlying cell state reprogramming, it is limited by sparsity of 3’ nuclear transcripts. We circumvented this limitation by also performing 3’ scRNA-seq from whole cells isolated from adults, D3-5, and Wk8 post transplantation using the 10x Genomics platform (**Figure 4A**). Together, these kinetic analyses capture the earliest molecular events in loss of basal identify, gain of luminal identity, and reveals the cell states present once the post-transplant mammary architecture has reached homeostasis.

**Figure 4.**
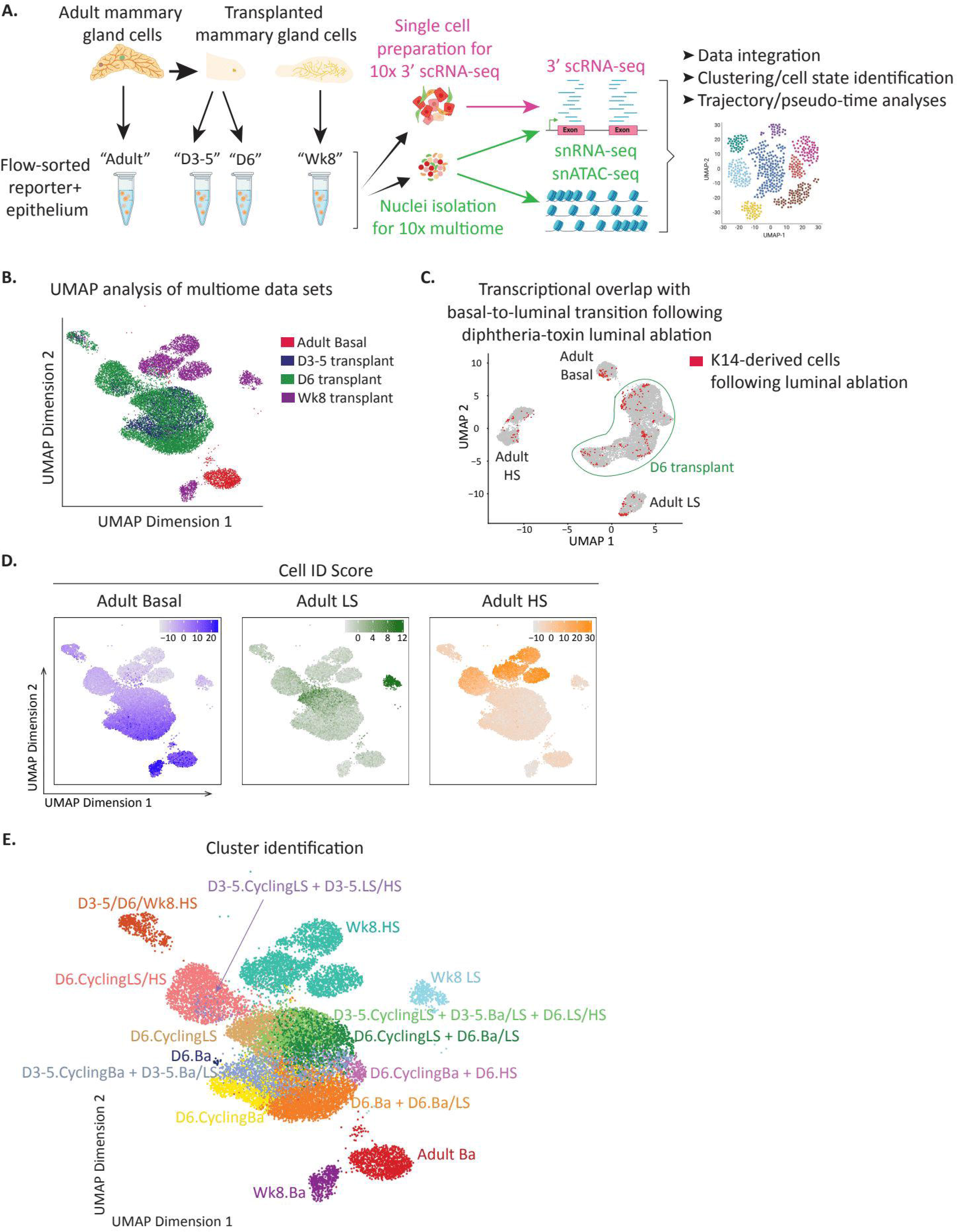
Multiome analysis reveals molecular dynamics of basal-to-luminal cell state transitioning. A. Schematic of mammary epithelial cells analyzed by multiome and 3’ scRNA-seq. “Adult” mammary epithelium isolated from K14-mCl; K18-tdT dual-reporter mice. “D3-5”, “D6”, and “Wk8” mammary epithelium from K14-mCl+ basal cell transplant recipient mice isolated 3-5 days (pooled), 6 days, and 8 weeks post-transplant, respectively. Created with BioRender.com. B. UMAP plot of cells by analysis timepoint. C. Transcriptional overlap of basal-to-luminal transitioning cells following diphtheria toxin luminal cell ablation^33^ (red) overlayed on adult and D6 transplant K14-mCl; K18-tdT mammary epithelium (gray). D. Cell ID scores from adult basal (left), LS (middle), and HS (right) overlayed on D3-5, D6, and Wk8 transplant cells. Transplant populations indicated in (B). E. Cluster identification in transplanted cells determined by timepoint, relatedness to adult cell ID signatures, and fraction of cells cycling.

We first focused our analyses on the multiome datasets. We leveraged UMAP analysis of combined ATAC and RNA datasets to subcluster and visualize cell populations from each timepoint (**Figure 4B**, see also Methods). We integrated data from different time point samples in order to address batch effects and employed the Seurat Integration protocol (see Methods). This revealed that, even as early as 3 days post-transplant, all transplanted cells (**Figure 4B**, blue and green) had epigenetically reprogrammed away from homeostatic adult basal cells (**Figure 4B**, red) demonstrating that transplantation rapidly enables basal cells to reprogram to a hybrid cell state. This rapid reprogramming is not an artifact of transplantation but rather represents a more general strategy by which fully differentiated mammary basal cells can repair tissue damage. We base this conclusion on our comparison of the single cell transcriptome data obtained above with one obtained from basal cells that transitioned to luminal *in situ* following diphtheria toxin-induced luminal cell ablation^33^. The overwhelming majority of *in situ* transitioning cells coincided with those identified in our D6 transplanted transitioning cells (**Figure 4C**). This demonstrates that basal-to-luminal conversion occurring in response to locally induced tissue damage and transplantation involve similar molecular mechanisms. The few *in situ* transitioning cells that overlapped with adult Basal, LS, and HS populations may have resulted from leaky lineage labelling, FAC sorting contamination, or true representation of cells that rapidly acquired luminal cell states in that context.

To closely track the molecular processes associated with loss of basal and gain of luminal cell states, we first developed cell ID scores for adult basal, LS, and HS based on our previously published snATAC-seq and 3’ scRNA-seq datasets^3, 44^ (**Figure S4A-C**, see methods) and compared these with cells undergoing basal-to-luminal conversion. Higher ID scores represent greater similarity to the respective adult population. Transcriptomic analysis revealed that cells in various stages of basal-to-luminal conversion did not bin into precise cell types as the Wk8 populations did, but rather the transitioning cells acquire mixed signatures associated by adult basal, adult LS, or adult HS cells (**Figure 4D**). This shows that our approach identified transitioning cells as those that exhibit a hybrid cell phenotype.

Considering we observed that the total cell number recovered in recipient mouse mammary fat pads increases significantly during 3-6 days post-transplant (**Figure 3E**), we evaluated proliferation within transitioning cell populations. Indeed, higher fractions of cycling cells, determined by *Mki67* RNA expression, was observed in certain subpopulations (**Figure S4D**). Therefore, we annotated subpopulations of transitioning cells based on the analysis timepoint, the adult population(s) they resemble most closely (taking into account that many transitioning cells co-express features from multiple adult cell types), and whether or not the population contained >10% *Mki67*+ cycling cells. For example, “D6.Ba + D6.Ba/LS” indicates a mix of D6 cells expressing basal-like signatures and D6 cells co-expressing basal-like and LS-like signatures, “D6.CyclingLS” indicates D6 cells with >10% *Mki67* expressing cells with LS-like signatures, and so on. Taken together, we identified a total of 15 clusters defined by time point, cell ID scores, and cycling status (**Figure 4E, Figure S4E-F**). We have generated a publicly-available shiny app to interrogate all of our multiome data sets, including transcript expression, chromatin accessibility, and programs linked to cluster transitions: https://tnbcworkbench.org/shiny/ShinyArchRUiO/.

### Adult basal cells de-differentiate to an earlier developmental state while transitioning to a luminal fate

The analyses above identified clusters within transitioning cells that group based on molecular signatures from multiple cell types, but not the potential routes by which basal cells transition to luminal cell fates. To unravel the molecular routes that enable changes in cell states following transplantation, we performed pseudotime analysis using Monocle to organize cells into a trajectory through time based on the epigenetic and transcriptional relatedness of individual cells. Starting with adult basal cells as the root cluster, this analysis identified 12 transitions spanning the diverse cell clusters that arise following transplantation (“**T1 – T12**”; **Figure 5A-B**). Of note, our pseudotime identified two independent transition routes away from adult basal: T1 and T11. T1 notably transitions to a hybrid cluster that generates multiple other cell states (e.g., through T2, T3, T4, T5) while T11 leads to a confined transition route through T7 that ends with a single cluster. Further, the trajectory analysis enables identification of genes (**Figure S5A**) and chromatin regions (**Figure S5B**) that reflect the orchestrated progression of cell states post-transplantation, contributing to our understanding of the underlying molecular dynamics and potential programs that enable cell fate switching.

**Figure 5.**
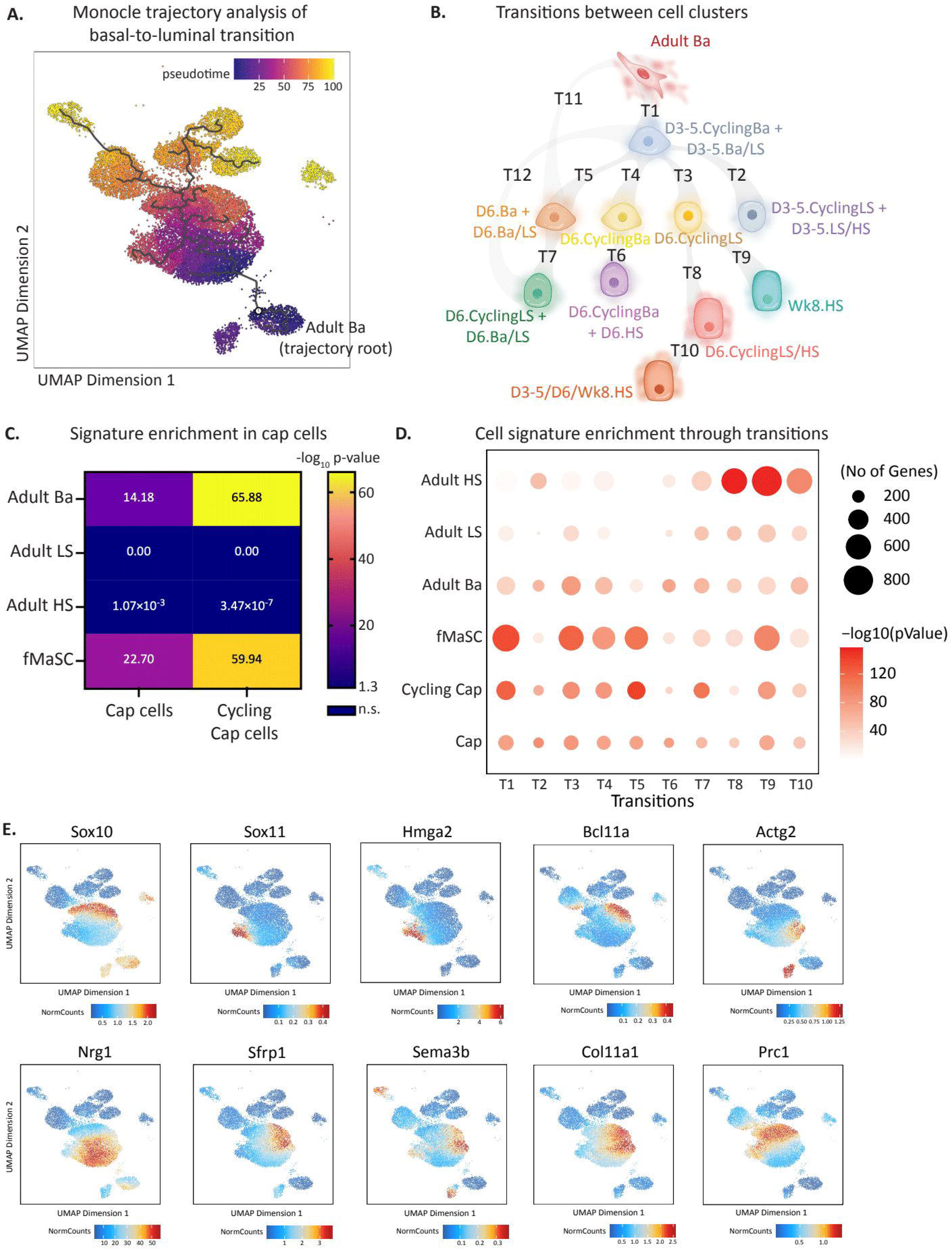
Adult basal cells de-differentiate to an earlier developmental state while transitioning to a luminal fate. A-B. UMAP plot (A) and schematic (B) of Monocle trajectory analysis starting with adult basal cells. The trajectory reveals 12 transitions through clusters that arise following basal cell transplantation and eventual mammary gland reconstitution. C. Cap cells, non-cycling and cycling, cell identity scores overlayed on fMaScs and adult basal (Ba), LS, and HS cells ID scores. P-value reflects the extent of similarity between ID scores. Graph displays −log_10_ P-value, so significance is ≥ 1.3 (−log_10_ 0.05 = 1.3). D. Bubble plot of cell signatures from adult HS, adult LS, adult basal, fMaSCs, cycling cap, and cap cells enriched in transitions identified in the Monocle trajectory analysis. E. Expression of fMaSC-enriched genes^3^ in transitioning cells.

Interestingly, the observation that cells in transition co-express molecular signatures from multiple cell types was reminiscent of what we previously observed to occur during embryonic mammogenesis. We and others previously showed that fMaSCs co-express genes typically expressed in basal and luminal cells, such as co-expression of basal and luminal keratins, K8 and K14^6, 27^, or lineage determining transcription factors for both basal and luminal lineages, Elf5 and Trp63^3^. Distinct lineages are evident shortly after birth, and final adult lineage molecular states are attained following puberty. During puberty the rudimentary mammary tree dramatically expands to give rise to a highly branched epithelial duct network filling the mammary fat pad. This process is orchestrated by hormonal cues that trigger terminal end bud (TEB) formation followed by ductal growth and elongation. TEB’s are specialized structures comprised of an inner multi-cellular body cell layer and an outer single-cell layer of cap cells. Cap cells, which are present through puberty, resemble adult basal cells but have been proposed to contribute to both basal and luminal lineages^8, 9^, thereby serving as a type of puberty-specific mammary stem cell subpopulation.

Given the plasticity observed in fMaSCs, the putative plasticity of cap cells, and the contribution of both in normal mammary gland development, we hypothesized that basal-to-luminal cell state change may proceed through a stem-like state resembling fMaSCs or perhaps cap cells. To address this possibility, we assessed the relatedness of the transitioning cell signatures to our previously derived fMaSC molecular signature. We also derived a molecular signature for cap cells by performing 3’ scRNA-seq analysis on mammary epithelial cells isolated from 4-5 week old female mice in puberty, as this developmental stage is linked to TEB elongation^48, 49^. We identified cap cells as a group of cells that were molecularly related to basal cells but distinguishable as a side population protruding from the basal cell cluster obtained from pubertal females (**Figure S5C**). As previously reported^7^, a subcluster of cap cells were proliferating (**Figure S5C-D**). Focusing on the non-cycling cap cell population to identify unique transcriptomic signatures that are not driven by proliferation-related programs, we observed significant gene expression differences between cap cells and age-matched pubertal basal cells (**Figure S5E**). Cap cells were enriched for epithelial-mesenchymal transition, translation, biosynthesis, and several gene networks linked to cellular metabolism as compared to non-cap basal cells from the same mice in puberty (**Figure S5F**). Of interest, when we compared cap cells against other mammary epithelial cell types, our data revealed that cap cells and cycling cap cells closely resembled fMaSCs (p < 1.82×10^-23^ and p < 1.15×10^-60^, respectively) and adult basal cells but not LS or HS cells, supporting the observation that they represent a stage in development between bipotent fetal cells and the homoeostatic adult basal population (**Figure 5C**). We have generated a publicly-available shiny app to interrogate our pubertal and cap cell scRNA-seq data and to be able compare it to adult populations: https://tnbcworkbench.org/shiny/shinyApp_cap/.

To determine if cells undergoing basal-to-luminal transition reprogram to an fMaSC-like or cap cell-like state, we overlayed the fMaSC and cap cell signatures on transitioning cells, focusing on T1 – T10 as these captured more precise transitions between closely-related clusters; T11 and T12, in contrast, represent transitions between more distant clusters (**Figure 5B**). This analysis revealed that transitioning cells, particularly those in T1, T3, T4, and T5, highly resemble molecular features comprising the fMaSC cell signature, with T1 being the most significantly fMaSC-enriched transitional state (**Figure 5D**). Further, we observed greater enrichment in the cap cell signature as compared to adult cell signatures in transitions T1 and T5, as well as a strong enrichment in the cycling cap cell signature in transitions T1, T5, and T7. This likely reflects both cap programs and cycling gene networks.

Considering the striking enrichment of the fMaSC signature in several transitional states, we analyzed which fMaSC-associated genes and/or programs are re-expressed during basal-to-luminal transition. We observed examples of characteristic fMaSC transcription factors^3^, including *Sox10*, *Sox11, Hmga2,* and *Bcl11a,* significantly upregulated in transitioning cells as compared to adult populations (**Figure 5E**). Cell-cell communication and extracellular matrix genes linked to the fMaSC state, such as *Nrg1, Sfrp1, Sema3b,* and *Col11a1* were also enriched in transitioning cells as well as cell motility (*Actg2*) and cell cycle (*Prc1*) genes (**Figure 5E**). Additionally, we observed the re-expression of previously reported postnatal chromatin modifiers, *Arid1a* and *Top2a^3^*, in transitioning cells but not in adult populations (**Figure S5G**). Notably, our analysis provided a continuous temporal view of these markers in transitioning cells, revealing some fMaSC genes appeared in the earliest transitioning clusters, such as *Nrg1, Sox11, Hmga2*, and *Actg2*, while others appeared later, such as *Sox10, Bcl11a*, and *Prc1*. While many of these genes were silenced by 8 weeks post-transplant, some, such as *Actg2* and *Sema3b*, maintained expression in Wk8 basal cells (**Figure 5E**). The putative stem cell markers *Tspan8*^45^ and *Procr*^28^ were not expressed in the adult basal lineage or in early transitioning cells, but rather expressed in later transitioning LS-like and HS-like cells, suggesting that the earliest steps in acquired plasticity does not reflect these genes previously reported to identify mammary stem cells (**Figure S5H**). Further, we did not observe the emergence of *Epcam*^Hi^/*Itgb1*^Hi^ co-expressing cells as previously reported in basal-to-luminal transition^33^, although we did observe potent induction of *Itgb1* in early transitioning cells (**Figure S5I**). Finally, we performed gene ontology (GO) analysis comparing early transitioning cells (T1) with fMaSCs and cap cells (**Figure S5J-K**). Transitioning cells exhibited chromatin reorganization and DNA replication similar to fMaSCs, and were enriched in translation and cell adhesion processes akin to cap cells. Taken together, these data suggest that cell transitions observed in T1, T3, T4, and T5 occur through de-differentiation into molecular states that resemble fMaSCs and also share features of pubertal cap cells.

### Loss of basal identity and transition to luminal state depends on Notch and BMP

To gain insight into the mechanims involved in the earliest steps of basal-to-luminal cell state transition, we identified genes in T1 with altered RNA expression as well as chromatin state as compared to adult basal (**Figure 6A**). As expected, transitioning cells downregulated the basal cell regulator *Trp63*. We unexpectedly observed altered expression of multiple cell signaling pathways including JAK/STAT, NF-κB, AP-1, and BMP. Gene Ontology analysis of pathways enriched in T1 revealed upregulation of cell cycle, DNA damage response, and protein translation (**Figure 6B and 6C, Figure S6A**), all of which are in line with the high proliferation we observed in these early time points as transitioning cells regenerate the mammary gland (**Figure 3E**). Enrichment of metabolic pathways including TCA cycle, respiratory electron transport, serine synthesis, and lipid metabolism could be attributed to the lack of blood vessels shortly after transplant, or triggered by the experimental manipulation.

**Figure 6.**
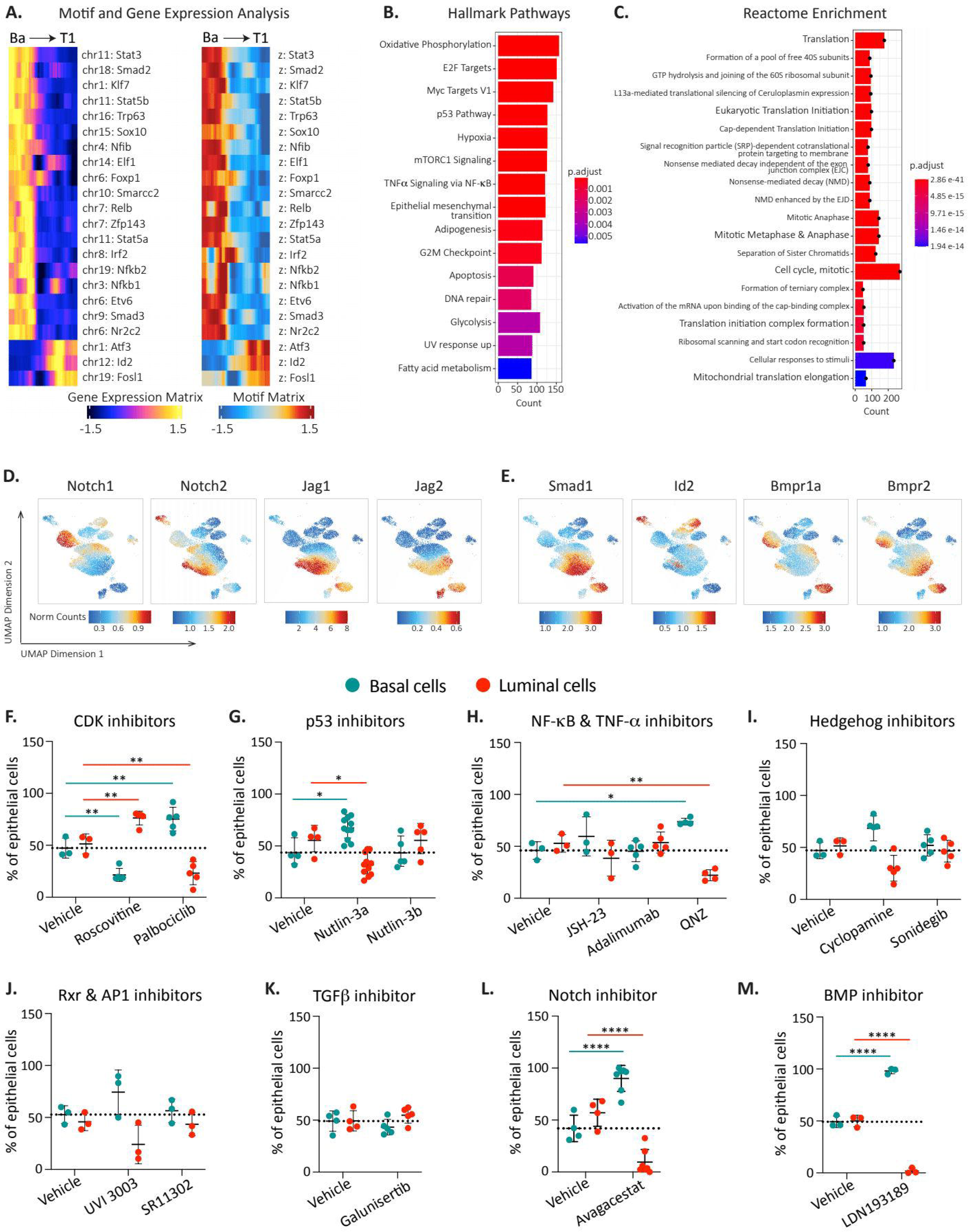
Loss of basal identity and transition to luminal state depends on Notch, BMP and Retinoid X Receptors. A. Motif and gene expression correlation analysis in T1 transition. B-C. Functional enrichment analysis on T1 transition for upregulated hallmark enrichment (B) and upregulated reactome enrichment (C). D. UMAP of selected Notch-related genes including Notch1, Notch2, Jag1, and Jag2. E. UMAP of selected BMP-related genes including Smad1, Id2, Bmpr1a, and Bmpr2. F-M. In vitro screening of inhibitors modulating basal-to-luminal transition. K14-mCl+ basal cells were sorted and cultured in Matrigel to form organoids. Cells were treated with various inhibitors for 8 days and analyzed via flow cytometry to assess the basal-to-luminal cell ratios. The percentage of basal and luminal cells among total epithelial cells is shown in the presence of inhibitors targeting: (F) CDK, (G) p53, (H) NF-κB and TNF-α, (I) Hedgehog, (J) Rxr and AP1, (K) TGF-β, (L) Notch, and (M) BMP pathways. Dotted lines indicate the percentage of basal cells in DMSO-treated vehicle controls for each condition. n = 3-11 biological replicates per condition. Data are represented as mean ± SD. Two-way ANOVA with Sidak’s multiple comparison test; * p ≤ 0.05; ** p ≤ 0.01; **** p ≤ 0.0001.

We next investigated the involvement of specific pathways implicated in normal fetal mammary development within transitioning cells. As expected, these cells were in a proliferative state, exhibiting time-dependent expression of various cyclins (**Figure S6B**). Interestingly, while we did not detect *Trp53* expression in transitioning cells, *Mdm2*, a known negative regulator of p53, was present (**Figure S6C**), suggesting downregulation of the p53 pathway as occurs in early embryonic cells^50, 51^. Additionally, we observed upregulation of several Hedgehog pathway components, particularly *Gli3* (**Figure S6D**). Notably, *Rxra* from the retinoid X receptor family was exclusively upregulated in a specific subset of transitioning cells, yet absent in adult epithelial cells (**Figure S6E**). The AP-1 pathway was also active, marked by high expression of *Fos* and its ligand (**Figure S6F**). In line with previous studies showing that TNF-α secreted by luminal cells restricts basal cell multipotency^33^, we found that *Tnf* was absent in transitioning cells and expressed only in alveolar luminal secretory cells (**Figure S6G**). Furthermore, NF-κB and its downstream targets were activated in a temporally specific manner (**Figure S6H**). Since earlier reports suggested TGFβ involvement in basal-to-luminal transitions during puberty^52^, we examined TGFβ and TGFβR expression. While TGFβ1 was not expressed in transitioning cells, TGFβ2 was upregulated in a specific subpopulation (**Figure S6I**). Interestingly, multiple Notch family members, both receptors and ligands, were upregulated in transitioning cells. *Notch2*, *Jag1*, and *Jag2* were expressed early post-transplantation, while *Notch1* and *Notch3* were expressed as cells began acquiring luminal characteristics (**Figure 6D**, **Figure S6J**). Similarly, BMP signaling members were progressively upregulated during basal-to-luminal transition (**Figure 6E**, **Figure S6K**).

To test if pathways identified by bioinformatic analyses functionally contribute to basal-to-luminal transition, we isolated K14-mCl+ basal cells from K14-mCl+; K18-tdT+ adult mammary glands and cultured them as organoids for 8 days in the presence of various signaling pathway inhibitors. Interestingly, Roscovitine, a CDK2/5 inhibitor, promoted basal-to-luminal conversion, whereas Palbociclib, a CDK4/6 inhibitor, blocked this transition (**Figure 6F**). Among the p53 modulators tested, Nutlin-3a, which activates p53 by inhibiting Mdm2, significantly inhibited basal-to-luminal conversion, while Nutlin-3b, an inactive enantiomer, served as a control (**Figure 6G**). Unexpectedly, while JSH-23 (an NF-κB inhibitor) and Adalimumab (a TNF-α monoclonal antibody) had no effect on mammosphere formation, QNZ, an inhibitor of both NF-κB and TNF-α, significantly reduced basal-to-luminal conversion (**Figure 6H**). We observed no impact on basal-to-luminal transition from Hedgehog, Rxr, or AP-1 pathway inhibitors (**Figure 6I-J**), and contrary to previous findings demonstrating the TGFβ inhibitor Galunisertib^52^ blocks cap cell-to-luminal transition, we did not observe any impact on basal-to-luminal conversion in adult basal cells (**Figure 6K**). It is possible that the soluble ligands normally activating these pathways are secreted by stromal or luminal populations, making inhibition in this context less critical for basal plasticity. Notably, the Notch inhibitor Avagacestat completely suppressed basal-to-luminal conversion, with almost no luminal cells observed in culture after 8 days of treatment (**Figure 6L**). Similarly striking, the BMP inhibitor LDN193189 effectively blocked basal-to-luminal conversion, with nearly all cells remaining in the basal state in its presence, highlighting a previously unappreciated pathway critical for basal cell plasticity (**Figure 6M**).

## Discussion

Injury-induced cell state transitions contribute to injury resolution and reacquisition of tissue homeostasis in several organs, including lung^29^, skin^30–32^, and intestine^53^. Recent studies in the mammary gland demonstrated that disruption of the epithelium by either diphtheria toxin-induced luminal ablation^33^ or exposure to DNA-damaging agents^34^ leads to basal-to-luminal cell state changes to effect tissue repair. However, the rate and mechanisms at which acquired plasticity arises remain unclear.

Here we illuminate the molecular framework by which fully differentiated adult mammary basal cells change to a luminal state. We used new mouse models in which genetically-engineered cell-state reporters enable tracking of dynamic cellular reprogramming that occurs during mammary gland reconstitution. The basal-luminal transition occurred very rapidly, initiating within days following transplantation which may be facilitated by the similarity between adult basal cell and bipotent fetal mammary stem cell epigenomes^44^. The earliest changes we observed revealed that basal cells rapidly underwent global epigenomic and transcriptomic reprogramming toward a basal-luminal hybrid phenotype overlapping with early developmental states. We suggest that the disruption of tissue structure unleashes an inherent potential of basal cells to undergo this remarkably rapid cell-state conversion.

While we cannot state what proportion of basal cells undergo this rapid cell state reprogramming, our observations indicate that it does not require invoking a rare, reserve basal stem cell pool. Rather, based on the speed of transition, the totality of transplanted cells that no longer molecularly resemble basal cells, and the short time required to regenerate a full and functioning mammary gland from low transplanted cell numbers, our findings strongly suggest that a rare, reserve basal stem cell pool is not responsible for mammary gland reconstitution. Consistent with this idea, we found no evidence of *Tspan8*^45^ or *Procr*^28^, two reported putative mammary stem cell markers, being expressed in adult basal lineage or early transitioning cells (**Figure S5G**). Rather, early transitioning cells upregulated fMaSC-associated genes, such as *Hmga2* and *Sox11* (**Figure 5E**) and further upregulated *Krt14* expression (data not shown). Induction of *Tspan8* and *Procr* expression was observed in cells becoming more luminal-like (**Figure S5G**). Our study corroborates previous reports supporting a single molecular cluster of basal cells by providing functional evidence that most, and possibly the majority, of basal cells may serve as facultative stem cells.

The transition from a basal to luminal state is likely complex, not the result of a single signaling cascade, transcription factor, or epigenetic modulator. The apparent co-expression of fetal and cap cell signatures suggests that individual cells do not directly transdifferentiate to luminal, but rather partially de-differentiate and then gain expression of differentiated luminal specifying genes. As we observed expression of the embryonic cell state regulator *Sox11* at early time points in a subset of cells (**Figure 5D-E**), we infer an early step involves progression through a fetal-like intermediate. Microenvironmental cells, including immune cells, are likely not required for this transition, as evidenced by extensive fate-switching in vitro following basal cell purification and relatively equal reconstitution frequencies in immune deficient and immune competent recipient mice (Q.V.M. and N.K.L., unpublished observations).

Blanpain and colleagues also demonstrated that fate-switching following diphtheria toxin-induced luminal ablation occurs through a CD29 (Itgb1)^Hi^; EpCAM^Hi^ intermediate requiring p21 and CDK1-linked cell proliferation and signaling pathways driven by TNF, PDGF, Notch, Wnt, and ErbB^33^. In contrast, Macara and colleagues reported that chemotherapy-induced luminal cell death leads to cap cell-derived basal-to-luminal interconversion, a process that is proposed to be independent of proliferation, Wnt, and Notch, but requires TGFβ, Jak2/Stat3, PI3K, MEK, and ROCK signaling and activation of a stromal inflammasome response^34^. Our epigenetic and transcriptomic data point to several molecular pathways that are critical for acquiring a hybrid cell state resembling that observed after luminal cell ablation^33^ (**Figure 4C**). Some of these corroborate previous studies, including a critical role for Notch signaling in the early stages of basal state downregulation and conversion to a luminal fate^27, 54^. Additionally, we observed an induction of *Itgb1* early in transition and *EpCAM* in later transition stages (**Figure S5I**) and demonstrate a critical role for CDK4/6- linked cell proliferation. However, our molecular and functional analyses suggest that TNF-α alone is dispensable for the transition (**Figure 6J**). Further, our studies highlight signaling axes that were not previously identified in basal-to-luminal transition such as Bone Morphogenic Protein (BMP), a class of potent growth factors critical for development, and NF-κB together with TNF-α, typically linked to stress response and inflammation (**Figure 6J, M**).

While our data suggest that basal cells rapidly acquire an epigenetic and transcriptomic transitional, hybrid state after transplantation, in order to generate an entirely new mammary gland, some of these cells must revert back to a differentiated, adult basal state while others proceed to form the luminal cell compartment. It is possible this could be the result of cell position and re-establishment of the basement membrane, such as occurs during development and in epithelial differentiation^55, 56^. Further, it remains unclear if a single basal-derived hybrid cell can give rise to both basal and luminal descendants in the reconstituted mammary gland, or if some hybrid cells are fated toward basal while others are fated toward luminal. Resolving this issue will require use of evolving barcoding approaches in future studies^57, 58^.

Since cancer has been referred to as the ever-healing wound that never heals^59^, it is tempting to speculate that the molecular programs involved in mammary basal-luminal cell fate switching also contribute to breast cancer initiation or the genesis of intratumoral heterogeneity. Indeed, hybrid cell states resembling those described here have been identified as intermediates in diverse cancers, and we and others have noted embryonic-like states in breast and other cancers^3, 6, 60–62^. Although diphtheria toxin-induced luminal ablation and basal cell transplantation are artificial models for injury-associated repair, the recent report showing that therapeutically relevant DNA-damaging agents cause basal-to-luminal switching^34^ supports the importance for understanding the immediate and long-term impact of mammary epithelial cell fate switching. It currently remains unknown if fate switching generates cells with enduring molecular alterations and if those cells could contribute to the growing challenge of second-primary breast cancer in breast cancer survivors^63, 64^. In this regard, we note that the basal cells analyzed 8-weeks post transplantation are epigenetically distinct from those used for transplantation (**Figure 4E**), suggesting that the epigenetic reprogramming linked to de-differentiation is not fully resolved even after the gland functionally appears homeostatic. Additionally, our study and previous ones^33, 34^ were performed in young, healthy mice and therefore may not capture the cell state changes linked to aging or menopause. Considering breast cancer incidence is highest in older, post-menopausal women and further linked to obesity, alcohol consumption, smoking, and premature weaning^65^, it will be important to investigate whether these risk factors link to increased cancer incidence by lowering the bar for cell-state instability. The new mouse model described here should provide a valuable tool for addressing such questions.

### Limitations of the study

We generated new mouse models that enabled us to elucidate the timing and molecular pathways by which pure adult basal cells regenerate an entire functional mammary gland within eight weeks of transplantation by re-isolating transitioning cells for single-cell epigenetics and transcriptomics. This technically challenging experimental design would not have been possible using in situ models with current technologies. However, by removing basal cells from their normal niche, we altered paracrine and juxtracrine signals between basal cells in contact, or from luminal cells, niche cells, and extracellular matrix components. We think this is not a serious limitation as our transplanted basal cell transcriptomes resemble those of transitioning basal cells following diphtheria toxin ablation of luminal cells^33^ (**Figure 4C**).

Finally, we cannot accurately estimate the fraction of basal cells that undergo cell state reprogramming. This would require introduction of barcodes into isolated basal cells prior to transplant. Any estimate we make may be affected by loss of basal cells we experienced at the earliest time points following transplantation (**Figure 3E**). It is possible there is cell loss during transplantation due to adhesion to plastic in the presence of Matrigel, and/or during dissociation and straining of the recipient gland. Conversely, cell loss could reflect the inability of some basal cells to survive transplantation, and this was not captured by our Annevin-V analysis (**Figure 3F**). Although our studies did not investigate mechanical losses, our data suggest that most, if not all, basal cells that survive were able to reprogram to a hybrid-like state. Therefore, while it remains possible that there is a rare, dedicated group of basal stem cells, our data are more consistent with the bulk of basal cells possessing the capacity to function as facultative stem cells when removed from their native cell:cell and cell:matrix/stroma constraints.

## Acknowledgments

We would like to thank Charles Perou (Univerity of North Carolina), Silvia Fre (Institut Curie), Joan Brugge (Harvard Medical School) for helpful discussions and scientific input. We thank A. Odelson, Y. Go, C. Ramos, L. Zamora, and A. Peterson for providing technical assistance with tissue and organoid dissociation. Instrumentation was supported by the Flow Cytometry Core Facility of the Salk Institute (RRID:SCR_014839) with funding from NIH-NCI CCSG: P30 CA01495, and Shared Instrumentation Grants S10-OD023689 (Aria Fusion cell sorter), and by the Waitt Advanced Biophotonics Core Facility of the Salk Institute with funding from NIH-NCI CCSG: P30 CA01495, NIH-NlA San Diego Nathan Shock Center P30 AG068635, and the Waitt Foundation. This work was supported in part by National Institutes of Health (NIH) R35 CA197687 (G.M.W), the Breast Cancer Research Foundation BCRF-23-168 (G.M.W.), the Hope Funds for Cancer Research Fellowship (N.K.L), American Cancer Society-Medical College of Wisconsin Cancer Center Institutional Research Grant (N.K.L.), Salk Women & Science Special Awards Initiative (N.K.L), the Wellcome-Leap Foundation (S.S.), NIH OT2-OD030544 (S.S.), NIH OT2-OD036435 (S.S.), Swiss National Foundation Mobility Fellowship P2ELP3_199760 (Q.V.M.), and the Catharina Foundation Fellowship (Q.V.M.). The funders had no role in the study design, data collection and analysis, or preparation of the manuscript.

## Author contributions

Conceptualization, G.M.W. and N.K.L.; methodology, Q.V.M, C.D., S.Sr., D.M., and N.K.L; bioinformatic analysis, Z.M., S.Sr., D.M., and K.M.; biological analysis, Q.V.M, C.D., and N.K.L; data curation, Q.V.M., Z.M., S.Sr., D.M., C.D., K.M., and N.K.L.; writing – original draft, N.K.L, G.M.W, Q.V.M.; figure generation, N.K.L. and Q.V.M; supervision, G.M.W, N.K.L, Q.V.M, and S.Su.; funding acquisition, G.M.W., N.K.L., and Q.V.M.

---

**Figure S1.**

A. Detailed design of mammary epithelial cell state indicator mice shown in Figure 1A. Basal cell reporters: Keratin14 (Krt14)–mClover (K14-mCl) and Krt14-tdTomato (K14-tdT). Luminal cell reporters: K8-tdT, K18-tdT, and K19-mCl. NLS (nuclear localization sequence) and 3NLS (three consecutive NLS) drive fluorophore nuclear localization for each cell state reporter. Self-cleaving 2A sequences separate each gene, indicated by triangles. When K14-mCl is crossed with K8-tdT or K18-tdT, K14;K8 or K14;K18 double-positive cells can be targeted for caspase-driven cell death by rapamycin-inducible cell killing systems.

B. Example gating scheme for identification and isolation of mammary epithelial cells by flow cytometry. Debris, cell doublets, and dead (Dapi+) or lineage+ (CD45+/Ter119+/CD31+) cells are initially excluded. Epithelial cells are then identified as EpCAM^Hi^; CD49f^Lo^ (luminal) or EpCAM^Lo;^ CD49f^Hi^ (basal).

C – G. Representative histograms of (C) K14-mCl expression, (D) K14-tdT expression, (E) K8-tdT expression, (F) K18-tdT expression, or (G) K19-mCl expression within total mammary epithelium (top row), basal cells (middle row), or luminal cells (bottom row).

H – I. Comparison of EpCAM^Lo^; CD49f^Hi^ basal cells with knock-in reporter expression (K14-tdT and K14-mCl) or previously published K14-promoter transgenic expression (K14-RFP). Representative flow plots (H) quantified in (I). n = 6-10 per group.

J. Representative z-stack images of K18-tdT; K14-mCl dual-reporter endogenous fluorescence in mammary buds from E18.5 female embryos. Yellow cells indicate dual expression of K18-tdT and K14-mCl. Scale bars: 100 μm; z-stacks: 10 μm (top) and 16 μm (bottom).

K. Representative dot plots of flow cytometry analysis of K18-tdT^Hi^ and K18-tdT^Lo^ luminal frequency within alveolar luminal secretory cell populations Itgb3+ (CD61+; top), CD49b+/Sca1- (middle), and cKit+ (bottom).

L. UMAP plot of mammary epithelial cells isolated from adult K18-tdT; K14-mCl and wildtype mice analyzed by 3’ scRNA-seq. Each data set is representative of cells pooled from n = 3 mice.

Data are represented as mean ± SD. One-way ANOVA with Tukey’s multiple comparison test; **** p ≤ 0.0001

**Figure S2.**

A. UMAP plot of mammary epithelial cells isolated from adult K18-tdT; K14-mCl analyzed by 3’ scRNA-seq. K18-tdT+/K14-mCl+ double-positive reporter cells (red) are overlayed on single-positive reporter cells (gray). Each data set is representative of cells pooled from n = 3 mice.

B. Expression of putative mammary stem cell genes *Tspan8* (top) and *Procr* (bottom) in K18-tdT+/K14-mCl+ double-reporter positive cells (left plots) or K14-mCl+/K18-tdT-, K18-tdT+/K14-mCl-single-reporter positive cells (right plots).

C. Expression of heat shock related proteins *Hspa1a* (left) and *Hspa1b* (right).

D. Heat shock transcriptomic signature overlayed on double-positive reporter cells (left) and single-positive reporter cells (right).

**Figure S3.**

A. Representative flow plots of reporter expression in transplant recipients at indicated time points. Each plot represents an independent biological replicate.

**Figure S4.**

A-C. Gene expression dot plots of adult basal (A), adult LS (B), and adult HS (C) associated genes.

D. Frequency of cells within each cluster expressing *Mki67.* Cycling cells are defined as *Mki67+*, and clusters that contain at least 10% *Mki67+* are labeled as cycling (indicated by red line).

E. Distribution of each cluster comprising Adult Basal, D3-5 transplant, D6 transplant, and Wk8 transplant.

F. Gene expression matrix with gene clusters significantly enriched in each cell cluster.

**Figure S5.**

A-B. RNA gene expression matrix (A) and ATAC peak matrix (B) depicting highly expressed genes or open chromatic regions, respectively, through T1 to T12 transitions identified by Monocle Trajectory analysis.

C. UMAP plots showing 3’ scRNA-seq analysis of K14-mCl; K18-tdT mammary epithelial cells isolated from adults (left; 2-4 month old) or pubertal (right; 4-5 weeks old). Non-cycling cap cells (teal) and cycling cap cells (blue) in the pubertal set are observed as a shoulder off of pubertal basal cells (red). Each data set is representative of cells pooled from n = 3 mice.

D. *Mki67* expression in pubertal K14-mCl; K18-tdT mammary epithelial cells.

E. Volcano plot of differentially expressed in non-cycling pubertal cap cells as compared to pubertal basal cells.

F. Gene Ontology analysis of non-cycling pubertal cap cell enriched genes vs. pubertal basal cells; Biological Processes (left) and Hallmark gene sets (right).

G. Expression of putative mammary stem cell genes *Tspan8* (left) and *Procr* (right).

H. Expression of postnatal chromatin modifiers *Arid1a* (left) and *Top2a* (right).

I. Expression of *Itgb1* (left) and *Epcam* (right).

J-K. Gene ontology analysis highlighting enriched Biological Processes found to co-occur in T1 and fMaSCs (J) or T1 and non-cycling cap cells (K).

**Figure S6.**

A. Gene Ontology enrichment analysis of Biological Processes upregulated (right column) or downregulated (left column) in T1 transition as compared to adult basal cells.

B-J. UMAP plots showing expression of selected genes involved in key biological pathways, including cyclins (B), p53 (C), Hedgehog (D), RXR (E), AP-1 (F), TNFα (G), NF-κB (H), TGFβ (I), Notch (J), and BMP (K).

## STAR Methods

### KEY RESOURCES TABLE

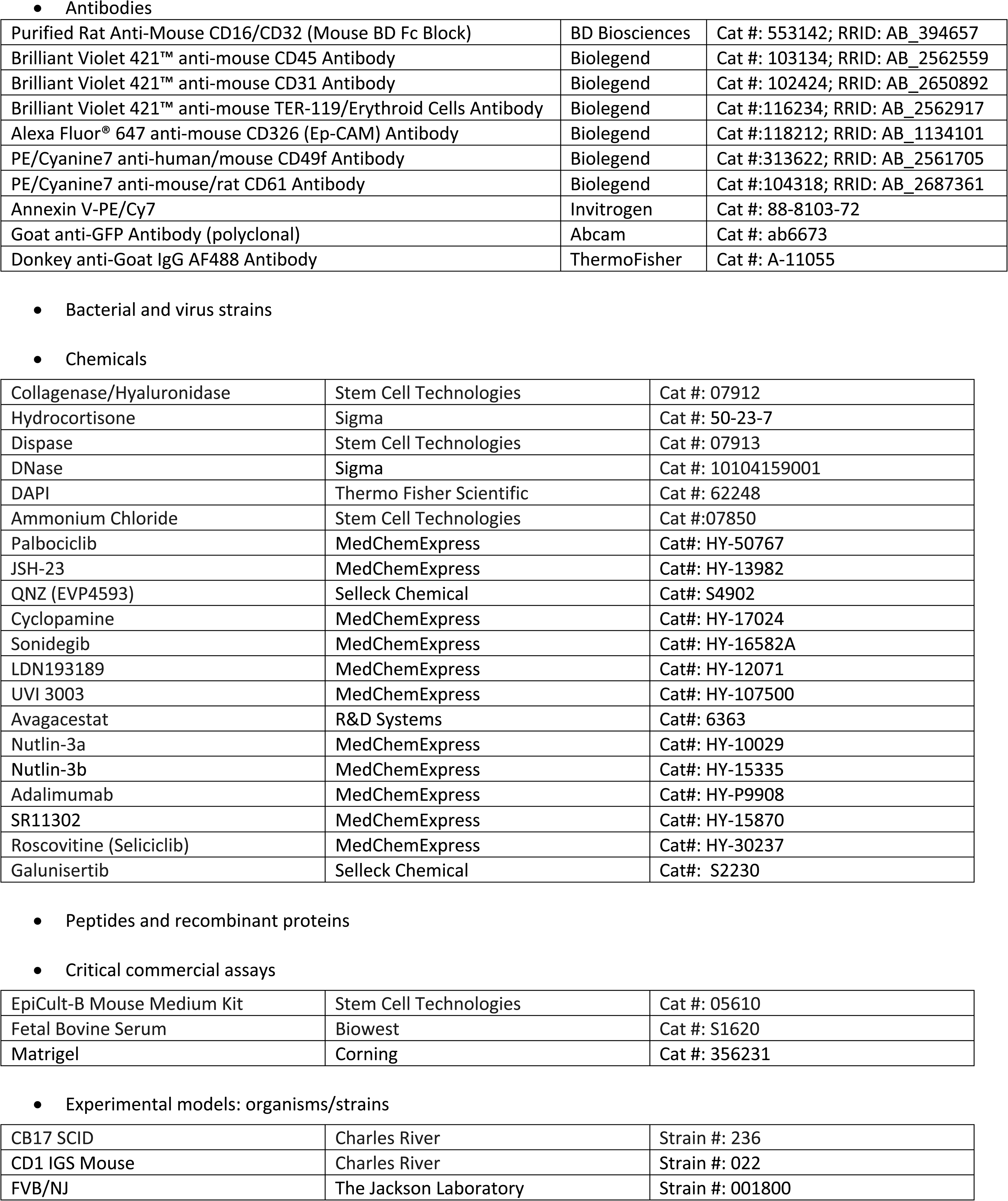
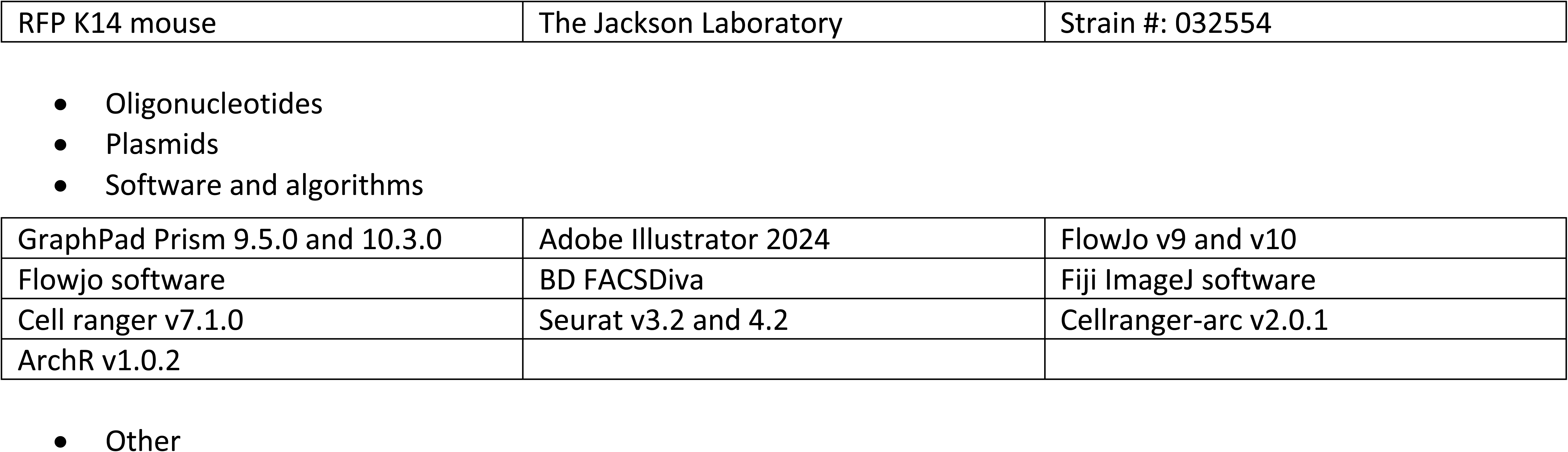

|  |  |  |
| --- | --- | --- |
| Purified Rat Anti-Mouse CD16/CD32 (Mouse BD Fc Block) | BD Biosciences | Cat #: 553142; RRID: AB_394657 |
| Brilliant Violet 421™ anti-mouse CD45 Antibody | Biolegend | Cat #: 103134; RRID: AB_2562559 |
| Brilliant Violet 421™ anti-mouse CD31 Antibody | Biolegend | Cat #: 102424; RRID: AB_2650892 |
| Brilliant Violet 421™ anti-mouse TER-119/Erythroid Cells Antibody | Biolegend | Cat #:116234; RRID: AB_2562917 |
| Alexa Fluor® 647 anti-mouse CD326 (Ep-CAM) Antibody | Biolegend | Cat #:118212; RRID: AB_1134101 |
| PE/Cyanine7 anti-human/mouse CD49f Antibody | Biolegend | Cat #:313622; RRID: AB_2561705 |
| PE/Cyanine7 anti-mouse/rat CD61 Antibody | Biolegend | Cat #:104318; RRID: AB_2687361 |
| Annexin V-PE/Cy7 | Invitrogen | Cat #: 88-8103-72 |
| Goat anti-GFP Antibody (polyclonal) | Abcam | Cat #: ab6673 |
| Donkey anti-Goat IgG AF488 Antibody | ThermoFisher | Cat #: A-11055 |
| Collagenase/Hyaluronidase | Stem Cell Technologies | Cat #: 07912 |
| Hydrocortisone | Sigma | Cat #: 50-23-7 |
| Dispase | Stem Cell Technologies | Cat #: 07913 |
| DNase | Sigma | Cat #: 10104159001 |
| DAPI | Thermo Fisher Scientific | Cat #: 62248 |
| Ammonium Chloride | Stem Cell Technologies | Cat #:07850 |
| Palbociclib | MedChemExpress | Cat#: HY-50767 |
| JSH-23 | MedChemExpress | Cat#: HY-13982 |
| QNZ (EVP4593) | Selleck Chemical | Cat#: S4902 |
| Cyclopamine | MedChemExpress | Cat#: HY-17024 |
| Sonidegib | MedChemExpress | Cat#: HY-16582A |
| LDN193189 | MedChemExpress | Cat#: HY-12071 |
| UVI 3003 | MedChemExpress | Cat#: HY-107500 |
| Avagacestat | R&D Systems | Cat#: 6363 |
| Nutlin-3a | MedChemExpress | Cat#: HY-10029 |
| Nutlin-3b | MedChemExpress | Cat#: HY-15335 |
| Adalimumab | MedChemExpress | Cat#: HY-P9908 |
| SR11302 | MedChemExpress | Cat#: HY-15870 |
| Roscovitine (Seliciclib) | MedChemExpress | Cat#: HY-30237 |
| Galunisertib | Selleck Chemical | Cat#: S2230 |
| EpiCult-B Mouse Medium Kit | Stem Cell Technologies | Cat #: 05610 |
| Fetal Bovine Serum | Biowest | Cat #: S1620 |
| Matrigel | Corning | Cat #: 356231 |
| - Experimental models: organisms/strains |  |  |
| CB17 SCID | Charles River | Strain #: 236 |
| CD1 IGS Mouse | Charles River | Strain #: 022 |
| FVB/NJ | The Jackson Laboratory | Strain #: 001800 |
| RFP K14 mouse | The Jackson Laboratory | Strain #: 032554 |
| GraphPad Prism 9.5.0 and 10.3.0 | Adobe Illustrator 2024 | FlowJo v9 and v10 |
| Flowjo software | BD FACSDiva | Fiji ImageJ software |
| Cell ranger v7.1.0 | Seurat v3.2 and 4.2 | Cellranger-arc v2.0.1 |
| ArchR v1.0.2 |  |  |

- Other

### RESOURCE AVAILABILITY

#### Lead contact

Further information and requests for resources and reagents should be directed to and will be fulfilled by the lead contact, N.K.L.; G.M.W. is now retired.

#### Materials availability

Mouse models (e.g. Keratin reporters) developed and used in this study are available upon request.

#### Data and code availability

- Sequencing, including single-cell RNA-sequencing and single-nucleus ATAC-sequencing paired with single-nucleus RNA-sequencing (multiome) will be deposited in public databases prior to publication.

## EXPERIMENTAL MODEL AND SUBJECT DETAILS

### Mouse Strains

Mice were cared for in accordance with the NIH guidelines in AAALAC-accredited facilities at the Salk Institute for Biological Studies under the approved Salk Institute Animal Care and Use Committee guidelines. All animals were house in a controlled environment, fed rodent chow *ad libitum,* and cared for by the Animal Research Department.

## METHOD DETAILS

### Mice

Keratin reporter mice were generated through CRISPR-mediated homologous recombination in R1 ES cells. Targeting vectors were designed to insert the indicated reporter sequences at the 3’ of the keratin loci to capture endogenous keratin expression. 2A sequences were used to separate the keratin and reporter sequences to minimize perturbation of endogenous keratin protein identity. Included on the targeting vectors were sgRNAs under U6 promoter control containing targeting sequences to cleave keratin loci near the site of targeting vector insertion. Also included on the targeting vectors were LoxP-flanked G418 resistance cassettes to mediate positive selection of targeted ES clones, and a 3’ DTA expression cassette to mediate negative selection of improperly targeted ES clones. R1 cells were targeted by electroporation of the indicated TV along with a vector encoding Cas9, with clones selected after 7-10 days, expanded individually, and screened by PCR. Positive clones that were PCR validated were further expanded for blastocyst injection to generate chimeric mice, with germline transmission used to establish keratin reporter animal strains.

For K19-mClover3 (e.g., clone “646”), 350 bp 5’ and 400 bp 3’ HA arms were utilized, along with the sgRNA CTTGCTTATTTCAGCCGATG to insert a 2A-mClover3NLS-2A-mClover3NLS-STOP sequence into frame of the 3’ of K19 just upstream of the stop site. The final products of this targeting are the expression of K19 and 2 mClover3(NLS) proteins from the K19 locus.

For K14-tdTomato (e.g., clone “658”), 635 bp 5’ and 601 bp 3’ HA arms were utilized, along with the sgRNA AGCAGTATCTGCGTCCACGC to insert a 2A-tdTomatoNLS-STOP sequence into frame of the 3’ of K14 just upstream of the stop site. The final products of this targeting are the expression of K14 and tdTomato(NLS) from the K14 locus.

For K14-mClover3NLS (e.g., clone “663”), 635 bp 5’ and 601 bp 3’ HA arms were utilized, along with the sgRNA AGCAGTATCTGCGTCCACGC to insert a 2A-FRB(NLS)-N-Tev-2A-Caspase3(Tev)-2A-mClover3(NLS)-2A-mClover3(NLS)-STOP sequence into frame of the 3’ of K14 just upstream of the stop site. The final products of this targeting are the expression of K14, FRB-fused N-terminal split Tev protease, a caspase3 with Tev cleavage sites, and 2 mClover3(NLS) proteins from the K19 locus.

For K8-tdTomato (e.g., clone “664”), 539 bp 5’ and 438 bp 3’ HA arms were utilized, along with the sgRNA GCAGCTGCTCAGGGCTCACG to insert a 2A-KFBP-C-Tev-2A-tdTomato(NLS)-STOP sequence into frame of the 3’ of K8 just upstream of the stop site. The final products of this targeting are the expression of K8, FKBP-fused C-terminal split protease, and tdTomato(NLS).

For K18-tdTomato (e.g., clone “747”), 436 bp 5’ and 499 bp 3’HA arms were utilized, along with the sgRNA CGCAAGCAGGAAGACTGCCA to insert a 2A-KFBP-iCaspase9-2A-tdTomato(NLS)-STOP sequence into frame of the 3’ of K18 just upstream of the stop site. The final products of this targeting are the expression of K18, FKBP-fused to iCaspase9, and tdTomato(NLS).

All keratin reporter mice are carried as heterozygous. Homozygous mice are viable but can develop skin lesions around 3-6 months of age. This is particularly evident in K14-reporter mice, which is likely due to reduced Keratin 14 gene expression from alleles harboring the knock-in fluorophore into the 3’ UTR region. CB17 SCID and CD-1 mice were purchased from Charles River. FVB/N-Tg(KRT14-RFP,-TK)4264Muth/J mice were purchased from The Jackson Laboratory. All mice were maintained in a pathogen-free animal facility at the Salk Institute for Biological Studies, housed on a 12-h light-dark cycle, at an ambient temperature of 23 °C, and 30–70% humidity. Water and food were provided *ad libitum*.

All mouse experiments were performed with approval from the Animal Resources Department at the Salk Institute for Biological Studies and Institutional Animal Care and Use Committee (IACUC), under the 2011-0005 protocol number.

### Embryonic reporter image analysis

Timed pregnancies were setup by crossing heterozygous adult K14-mCl; K18-tdT dual-reporter males with adult CD-1 females. The morning on the day a plug was found was designated as E0.5 and the plugged females were then single housed. E12.5, E14.5, and E16.5 embryos were retrieved for fluorescent imaging. E18.5 embryos were retrieved for mammary rudiment isolation for confocal imaging and/or flow cytometry analysis. Embryos were staged according to morphological criteria to confirm gestation period and sexing. Female embryos carrying both K14-mCl and K18-tdT reporters were used for experiments. For E18.5 studies, mammary rudiments were picked under a dissecting scope (Leica MZ16F Stereo). For imaging, rudiments were placed on top of a glass slide and directly imaged in a confocal microscope (Olympus FluoView FV3000 Confocal Microscope). Z-stacks were taken at 0.5 μm intervals.

### Whole mount mammary gland preparations and stainings

Mammary glands were processed and imaged following the FUnGI^43^ clearing protocol previously established with minor modifications. Briefly, mouse mammary glands were dissected and spread onto glass slides, then fixed in 4% methanol-free formaldehyde (ThermoFisher, 28908) for 45 minutes at 4°C on a shaker to preserve endogenous fluorescent proteins. After fixation, samples were rinsed in PBS with 0.1% Tween-20 for 10 minutes at 4°C on a shaker. The tissues were then detached from the slides and incubated in 8 ml of Wash Buffer (PBS, 0.1% Tween-20, 50 μg/ml ascorbic acid, 0.05 ng/ml L-glutathione reduced) on a roller mixer for 2 hours at 4°C. Blocking and permeabilization were achieved by incubating the tissues in 8 ml of Wash Buffer 1 (PBS, 0.2% Tween, 0.2% Triton, 0.02% SDS, 0.2% BSA, 50 μg/ml ascorbic acid, 0.05 ng/ml L-glutathione reduced) for 6 hours at 4°C. Mammary glands were then transferred to 1 ml of Wash Buffer 2 (WB2: PBS, 0.1% Triton, 0.02% SDS, 0.2% BSA, 50 μg/ml ascorbic acid, 0.05 ng/ml L-glutathione) containing a 1:200 dilution of primary antibody (goat anti-GFP, Abcam, ab6673) to detect mClover. While tdTomato fluorescence was preserved through fixation, mClover required immunolabeling. Samples were incubated for 120 hours at 4°C on a shaker, with daily rotation. Following primary antibody incubation, samples were washed three times in 5 ml of WB2 and left in WB2 overnight. The tissues were then incubated in 1 ml of WB2 containing a 1:500 dilution of secondary antibody (donkey anti-goat AF488, ThermoFisher, A-11055) and DAPI at 1:1000 dilution for 48 hours at 4°C on a shaker. After three additional washes in 5 ml of WB2 and an overnight incubation in WB2, mammary glands were transferred to 5 ml of FUnGI clearing solution (50% glycerol, 2.5 M fructose, 2.5 M urea, 10.6 mM Tris Base, 1 mM EDTA, 50 μg/ml ascorbic acid, 0.05 ng/ml L-glutathione) on an orbital mixer overnight at 4°C in the dark. Before imaging, samples were left at room temperature for at least 2 hours. Images were acquired using an inverted confocal laser-scanning microscope (Laser Scanning Confocal Light Microscopy Zeiss Airyscan 880-1) and processed using Fiji.

### Embryonic rudiments and mammary gland dissociation

E18.5 mammary rudiments (all #10 rudiments), adult (2-6 months old) nulliparous mammary glands (#4 abdominal mammary glands), and transplanted mammary glands were dissected, minced into small pieces and collected in dissociation media containing Epicult-B Basal medium (Stem Cell Technologies, 05610) supplemented with Epicult B supplement, fetal bovine serum (FBS, 5%, Peak Serum, PS-FB2), penicillin G and streptomycin (Pen/Strep, 1%, Gibco, 15140-122), amphotericin B (2.5 μg/ml, Gibco, 15290-026), hydrocortisone (4 μg, Sigma, 50-23-7), and collagenase/hyaluronidase (1650 U/550 U, Stem Cell Technologies, 07912). Samples were incubated at 37°C with shaking for 1.5 h for the fetal tissue, and 3 h for adult tissue. Samples were processed in sets of four to achieve consistent exposure of enzymes used to digest the tissues. Next, cells were manually dissociated by pipetting with dispase (5 U, Stem Cell Technologies, 07913) and DNase (100 μg, Sigma, 10104159001) solution during 30 s intervals for a total of 4 min, followed by 4 min incubation with trypsin (0.25%, GenClone, 25-510). Erythrocytes were then removed by incubation with ammonium chloride (0.6%, Stem Cell Technologies, 07800) for 2 min at room temperature (RT). Final suspensions were passed through a 40 μm cell strainer to remove aggregated cells. Last, cells were resuspended in Hank’s balanced salt solution with 2% FBS (HF buffer) containing Fc block (1 μg/ml, BD Biosciences, 553142) and DNase (10 μg), ready for flow cytometry staining.

### Single cell preparation and Fluorescence-Activated Cell Sorting (FACS)

Cells were stained at 4°C for 20 min with lineage markers including CD45-BV421 (Biolegend, 103134), Ter119-BV421 (Biolegend, 116234), and CD31-BV421 (Biolegend, 102424), epithelial markers including CD326 (EpCAM)-Alexa Fluor 647 (Biolegend, 118212), CD49f-APC/Cy7 (Biolegend, 313628), CD61-PE/Cy7 (Biolegend, 104318), CD49b-PE/Cy7 (Biolegend 103517), Sca-1-BV605 (Biolegend 108133), cKit-AP (Biolegend 105811), and Dapi to exclude dead cells. To analyze apoptotic cells, cells were further stained with the Annexin-V-PE/Cy7 apoptosis detection set (Invitrogen, 88-8103-72). Cells were sorted in a FACSAriaTM Fusion sorter. Single, live and lineage negative cells were selected, and gated for EpCAM^Hi^/CD49f^Lo^ for luminal, EpCAM^lo^/CD49f^Hi^ for basal, EpCAM^Hi^/CD49f^Lo^/CD61^+^ for alveolar luminal secretory, and EpCAM^Hi^/CD49f^Lo^/CD61^-^ for hormone sensing cells. Immediately after sorting, cells were manually counted and resuspended with HF for transplantation, or further *in vitro* experiments.

### 3D mammosphere forming assay

Sorted basal cells (500 cells) were seeded in ultra-low attachment 96 well plates (Corning, 3474) with 100 μl media containing Advanced DMEM/F-12 (Gibco, 12634-010), B27-Supplement (1X, Gibco, A35828-01), Glutamax (1X, Gibco, 35050-061), Pen/Strep (10,000 U/ml, Gibco, 15140-122), mEGF (10 ng/ml, Thermo Fisher Scientific, PMG8041), FGF (25 ng/ml, Corning, 354060), hydrocortisone (0.5 μg/ml, Sigma, 50-23-7), insulin (5 μg/ml, Sigma, I9278), and Matrigel (2%, Corning, 356231). Cells were cultured at 37°C in a humidified atmosphere at 7% CO_2_. On days 4, 8, and 12 post-seeding, 50 μl of fresh media was added on top of each well. For imaging, mammospheres were transferred at different time points into imaging chambers (Ibidi, 80826) and directly imaged. Images and z-stacks were acquired on an inverted confocal laser-scanning microscope (Laser Scanning Confocal Light Microscopy Zeiss Airyscan 880-1) and processed in Fiji. Z-stacks were taken at 1 μm intervals.

### In vivo mammary gland transplantation and analysis

Mammary gland fat pad clearing and subsequent cell transplantation were performed following established protocols^13^. Briefly, the 4th inguinal mammary glands (both left and right sides) of 21-day-old female CB17-SCID mice were surgically removed from the nipple to the lymph node, leaving epithelial-free cleared fat pads as transplantation sites. Sorted single basal cells, in varying numbers, were resuspended in 2-3 µl of HF buffer, mixed with 1 µl of Trypan blue (0.05%, Sigma, T8154), and 3-4 µl of Matrigel (Corning, 356231; the final mixture always contained 50% Matrigel). The cell suspensions were then transplanted using pre-cooled 0.3 cc, 31G ultrafine needle insulin syringes (BD, 328438). Skin flaps were sutured and stapled post-transplantation. Transplanted glands were excised at designated time points and dissociated into single cells for subsequent analysis as previously described.

### Sample preparation for single cell sequencing studies

All samples 10x 3’ scRNA-seq (v3.1), and 10x single cell Multiome ATAC + gene expression (v1.0) were prepared from freshly isolated cells and immediately subjected to GEM generation & Barcoding. Live, lineage-, reporter+ epithelial cells were isolated by FAC sorting.

For D3-5 post-transplant cells, basal cells (K14-mCl+/K18-tdT-) isolated from 8 (for D5), 10 (for D4), and 10 (for D3) adult female dual-reporter mice were transplanted into postnatal day 20-21 SCID mice on days 0, 1, and 2, respectively, then re-isolated from all recipient mice on day 5. Therefore, these samples captured cells from days 3, 4, and 5 post-transplant. Cells from recipient mice from each transplant day were pooled together at a 1:1:1 ratio prior to sequencing. Cell transplantation for D6 samples were performed on a single day, representing 12 biological replicate donor mice. Recipient cells isolated 6 days post-transplant were pooled together for sequencing. A similar approach was used for Wk8 samples representing 3 biological replicate donor mice.

For double-positive (K14-mCl+/K18-tdT+), #4 mammary glands from 3 dual-reporter adult females were dissociated for FAC sorting. For each mouse, double-positive epithelial cells were sorted as one population, and all K14-mCl+/K18-tdT- and K18-tdT+/K14-mCl-single-reporter positive cells were sorted as a pooled second population. All double-positive cells from all 3 biological replicates were then pooled together for sequencing. Single-reporter positive cells were pooled from 3 donors at a 1:1:1 (Basal:LS:HS) ratio prior to sequencing.

For 3’ scRNA-seq, cells were then pelleted, resuspended in PBS, and counted using trypan to exclude dead cells. For Multiome, cells were pelleted, nuclei isolated per 10x protocol, then counted using trypan as an indicator of nuclei. 9000 live cells or 9000 nuclei were loaded per gel bead sample for 3’ scRNA-seq or Multiome, respectively, with the exception of double-positive reporter cells, in which we were only able to load 1900 live cells for 3’ scRNA-seq. For each sample, cells or nuclei were loaded onto a microfluidic chip (10x Genomics Inc.) and library preparation with indexing was performed per the 10x manual. Final library quality and size were assessed by TapeStation (Agilent Biosystems) and Qubit (ThermoFisher) and libraries were pooled in equal molar ratio and sequenced on Illumina NovaSeq 6000.

### ScRNA-seq data processing

10x scRNA-seq fastq files were processed with cell ranger (v7.1.0) using default parameters. Count matrices were computed using *Read10X()* from the filtered barcode count matrix at each stage and was read into Seurat (v3.2), object for quality control and post processing. Then the expression matrix was read-depth normalized and log transformed with the *NormalizeData()*. Top 2000 most variable genes across all the single cells were used to run *ScaleData()* followed by PCA analysis. Filtering of the data was performed at cell and gene levels, to obtain reduced datasets, which were then used for downstream integration. Only the 37952 common features across all datasets were considered for all downstream analyses.

### Data filtering

For each sample identified above, we performed cell-level filtering by excluding cells based on the following quality criteria: (1) > 10% mitochondrial reads, (2) < 300 genes(counts), and (3) < 500 detected transcripts(features).

Additionally, we performed gene-level filtering by excluding genes that were expressed in less than 10 cells. To facilitate downstream integration, we only utilized the 2000 features/genes common across all datasets for further analysis.

### Data integration

Each sample was filtered, log normalized, and scaled prior to integration. A total of 2000 most variable features were detected within each dataset using the vst method implemented in Seurat. Subsequently, the normalized datasets were integrated using the *IntegrateData()* function of Seurat, on 2000 gene anchors identified via *FindIntegrationAnchors()* (*ndims = 2000*). The final integrated dataset considered for this analysis contained 32287 features across a total of 11343 cells from 2 samples.

### Data scaling and dimensionality reduction

Data scaling and dimensionality reduction were performed on the integrated dataset. We identified PC threshold by calculating where the principal components start to elbow and used *cumsum()* to determine point (*ndims = 19*) where the percent change in variation between the consecutive PCs is less than 0.1%. Subsequently, we performed *RunUMAP()*, *FindNeighbors(),* and *FindClusters()* using the Louvain algorithm with a *res = 0.5* to get meaningful clusters.

### Establishing the cellular cluster identity of the integrated dataset

Cluster identities were established using the following markers, basal (*Trp63*, *Krt5*, *Krt14*), alveolar luminal secretory (*Elf5*, *Ehf*, *Kit*), hormone sensing (*Esr1*, *Foxa1*, *Prlr*), and cycling (*Mki67*). Further the annotation was validated using *FindAllMarkers()* for each cluster with Wilcoxon rank sum test at least 25% expressed and Bonferroni-corrected p-values <0.05.

### Multiome analysis preprocessing

In this study, we performed a single nuclear multiome analysis using snRNA data from various time point samples (Adult, Day 3-5, Day 6, and Week 8). The raw fastq files were initially preprocessed with cellranger-arc (v2.0.1) to generate fragments and counts files for downstream analysis. Subsequently, the fragments files were processed using ArchR (v1.0.2) while the *filtered_feature_bc_count_matrix.h5* files underwent processing using Seurat (v4.2) pipelines.

### SnATAC-seq data quality Control

The sample fragment files were converted into ArchR Arrow files using the *createArrowFiles()* function and then imported into the ArchR project. To ensure data quality and reliability, we subjected the cells to rigorous filtering based on their signal-to-background ratio, which is measured by the TSS enrichment score. Only cells with TSS scores between 10 and 35 were retained for further analysis. Additionally, we filtered cells based on the number of unique nuclear fragments, setting a cutoff of unique fragments > 2000. After applying these quality control steps, we identified a total of 28,748 high-quality single nuclei. These cells were derived from different time point samples and were distributed as follows: 7,079 (71%) from the adult time point, 4,099 (87%) from Day 3-5, 13,011 (92%) from Day 6, and 4,559 (87%) from Week 8 (% indicates fraction of sequenced nuclei that passed quality control and were included in the analyses). The median number of high-quality ATAC fragments per cell ranged from 4,802 to 27,584.

### SnRNA-seq data quality control

In the subsequent steps of our analysis, we performed quality control on the snRNA data to ensure that only high-quality cells and genes were included in the analysis. For cells, we filtered out dead cells by setting a threshold for the number of features (genes) detected in each cell (feature counts > 750), and a range filter on the total counts per cell (between 1000 and 15000). Thus, only reliable and informative single nuclei were kept for further analysis.

### Normalization, scaling and data integration

Next, we integrated data from different time point samples in order to address batch effects and employed the Seurat Integration protocol. Each sample underwent filtering, log normalization, and scaling to prepare the data for integration. We applied read-depth normalization and log transformation using the *NormalizeData()* function. This step was crucial to make gene expression data comparable across cells and to enhance the effectiveness of subsequent analyses. To reduce the dimensionality of the dataset and focus on the most informative genes, we identified the top 2000 most variable genes across all single nuclei and scaled the data using the *ScaleData()* function and the variance stabilizing transformation (vst) method in Seurat. The scaling of data is crucial for the subsequent analysis since large variances in data will lead to biased results and therefore transforming the data to comparable scales can prevent this problem.

To ensure robust integration, we selected features that exhibited consistent variability across multiple datasets using the S*electIntegrationFeatures()* function. Next, we identified anchors, which are shared data points that serve as reference points for integrating the datasets, using the *FindIntegrationAnchors()* function. Anchors help establish meaningful connections between the different datasets and facilitate their combination. The actual integration of the datasets was then performed using the *IntegrateData()* function in Seurat. By incorporating the previously determined anchors, we successfully merged the data from all time point samples into a unified and integrated dataset. The final integrated dataset contained a total of 23,596 nuclei and encompassed 34,274 features (genes) across the four samples.

### PCA, clustering and annotation

Principal Component Analysis (PCA) is a dimensionality reduction method, used to reduce the dimensionality of large data sets while retaining as much information as possible. Principal components are new variables that are constructed as linear combinations or mixtures of the initial variables. These combinations are done in such a way that the principal components are uncorrelated and most of the information within the initial variables is squeezed or compressed into the first components. To determine an appropriate number of principal components (PCs) for subsequent analyses, we calculated the point where the percentage of explained variances and principal components started to show an elbow-shaped curve. We used the *cumsum()* function to identify the point (*ndims = 15*) where the percent change in variation between consecutive PCs dropped below 0.1%. This allowed us to retain the most significant PCs while reducing the dimensionality of the data. Subsequently, we performed *RunUMAP()*, *FindNeighbors()*, and *FindClusters()* using the Louvain algorithm with a resolution parameter (*res = 0.5*). These steps were instrumental in creating meaningful clusters of cells, facilitating the identification of distinct cell populations based on their gene expression profiles. Cluster identities were established using the following markers specific to murine mammary cell types namely, basal (*Trp63*, *Krt5*, *Krt14*), alveolar luminal secretory (*Elf5*, *Ehf*, *Kit*), hormone sensing (*Esr1*, *Foxa1*, *Prlr*) and cycling (*Mki67*).

### Filtering of multiplets for ATAC

In the next step of our analysis, we conducted barcode matching between snATAC and snRNA data to incorporate the gene expression values from the paired snATAC-seq + snRNA-seq multi-modal assay. This was achieved by adding the gene expression matrix to the ArchRProject using the *addGeneExpressionMatrix()* function.

To ensure the removal of doublets from our dataset, we did not utilize the built-in function *filterDoublets()* provided by ArchR. Traditionally, doublets detection relies on the hybrid signature generated from heterotypic doublets arising from different cell types. This may false positively remove cells with biologically hybrid status, and is not very effective in removing multiplets formed from cells of similar profile. Instead, we ultilized a novel read count-based method (AMULET^66^) for multiplet detection from snATACseq space of the multiome data. Uniquely, AMULET emurate regions with greater than two uniquely aligned reads across the genome to effectively detect multiplets. After filtering multiplets, we identified a final set of high-quality nuclei for further analysis. The number of retained nuclei from each time point sample were as follows: 5,006 adult nuclei, 2,547 from Day 3-5, 10,166 from Day 6, and 3,556 from Week 8. These represent the reliable and informative single nuclei that will be the basis for the subsequent analyses.

### Normalization and clustering of ATAC

Given the sparsity inherent in ATAC data due to numerous zero positions representing absence of binding information, traditional dimensionality reduction methods like PCA are less effective due to high inter-cell similarity at these zero positions. To address this, Latent Semantic Indexing (LSI), originally used in text analysis, has been adapted for ATAC-seq data normalization. LSI reduces the dimensionality of ATAC-seq count matrices while capturing biological variation by decomposing the matrix into "latent semantic" factors, revealing patterns of co-occurring peaks across samples. Here, we employed LSI for dimensionality reduction through *addIterativeLSI()* with *clusterParams* (*res = 0.2*, *sampleCells = 10,000*) and *varFeatures = 1,000* for normalization. To enhance batch correction, we incorporated the Harmony tool via *addHarmony()*, augmenting the LSI algorithm to yield stronger batch correction and embedding cells into a shared space grouping by cell type rather than dataset conditions. Graph-based clustering using *addClusters()* with parameters *resolution = 0.8* was performed in ArchR to achieve better separation and visualization. Subsequently, we introduced UMAPS for combined RNA and ATAC assays using *addUMAP()* to enable data visualization. Next the clusters were annotated based on snRNA clustering using *mapLabels()* function.

For focused analysis, we excluded adult LS and adult HS cells, as only adult basal cells were transplanted, while the remaining cells (D3-5, D6, and W8) were considered. Cluster annotation of matched snATAC-snRNA was achieved using a confusion matrix involving snATAC cluster IDs and snRNA labels.

### Pseudo bulk aggregate generation, peak calling, and identification of marker peaks

The prominent aspect of ATAC analysis involves peak calling and the identification of marker peaks in ATAC data. Given the sparse nature of ATAC data in the single-cell space, pseudo bulk replicates are employed as a strategy to perform peak calling. These pseudo bulk replicates play a pivotal role by aggregating individual cell data into a bulk-like representation, effectively reducing noise, and bolstering the statistical power of peak calling. ArchR generates multiple such pseudo-bulk samples for each designated cell grouping, aiming to group together similar single cells while not emphasizing the differences between them.

These cell groupings typically stem from individual clusters or collections of clusters that correspond to recognized cell types. To facilitate the process, the pseudo-bulk ATAC replicates were generated using the *addGroupCoverages()* function, where parameters such as *minCells = 750*, *maxCells = 1000*, and *minReplicates = 3* were employed. The underlying assumption is that these grouped single cells share sufficient similarity, thus enabling meaningful peak calling and downstream analyses on the aggregated data. This technique is particularly beneficial in cases where calling peaks on an individual cell basis is impractical due to data sparsity.

In ArchR, the process of detecting and merging overlapping peaks is carried out iteratively in a tiered manner using the *addReproduciblePeakSet()* function. This function utilizes the Macs2 tool^67^ for peak calling, leveraging its capabilities in identifying enriched regions, or peaks, within DNA sequences associated with distinct protein interactions like transcription factor binding or histone modifications. Macs2 employs statistical modeling to effectively handle background noise, enhancing peak calling accuracy and supporting insights into regulatory elements and functional genomics studies. The tool models the spacing between paired forward and reverse strand peaks, sliding a window across the genome to locate regions of enrichment that exhibit an M-fold higher read count compared to the background.

For *addReproduciblePeakSet()*, specific parameters such as *cutoff = 0.05* and *method = q* were employed, while minCells were determined similarly to the group coverages call. This was followed by adding the peak matrix to the ArchR project by *addPeakMatrix()* function. Marker peaks, which are peaks unique to specific cell groupings and offer insights into cluster- or cell type-specific biology, were identified using the *getMarkerFeatures()* function with parameter *useMatrix = PeakMatrix*. This process utilized default parameters as established by Wilcoxon pairwise comparisons and the resulting marker peaks were subsequently visualized using *plotMarkerHeatmap()* for better understanding and interpretation.

### Moaf enrichment, ChromVar deviations and CoAccessibility

Once a robust peak set is established, it becomes crucial to predict which transcription factors might be involved in the binding events that generate those accessible chromatin sites. This information aids in assessing marker peaks or differential peaks, enabling an understanding of whether these peak groups are enriched for specific transcription factor binding sites. For instance, we often observe an enrichment of lineage-defining transcription factors in cell type-specific accessible chromatin regions. To explore enriched motifs in peaks that are up or downregulated across various cell types, we incorporated motif annotations into the ArchR project using the *addMotifAnnotations()* function, with parameter *motifSet = cisbp* where CIS-BP is an online database of transcription factor (TF) binding specificities motif set employed for annotation^68^. Subsequently, we identified motif enrichments for marker peaks using the *peakAnnoEnrichment()* function, setting the parameter cutoff as *FDR ≤ 0.1* & *Log2FC ≥ 0.5*. To visually comprehend these motif enrichments across all cell groups, we utilized the *plotEnrichHeatmap()* function.

However, it is important to note that these motif enrichments are not calculated on a per-cell basis and they disregard the insertion sequence bias of the Tn5 transposase. To address this, we leveraged chromVAR^69^, an R package designed to predict the per-cell enrichment of TF activity from sparse chromatin accessibility data. By measuring deviations with respect to average background peaks (that was computed using the *addBgdPeaks()* function call) and generating bias-corrected z-scores for each motif, chromVAR provides a more nuanced view of per-cell TF activity. We computed per-cell deviations using the *addDeviationsMatrix*() function, creating a MotifMatrix that was then integrated into the ArchR project, thus accounting for individual cell differences and addressing sequence bias concerns.

Co-accessibility refers to the correlation in accessibility levels between two peaks across multiple single cells. This metric holds the potential to predict regulatory interactions and enhancer activity through peak-to-gene linkage analysis. To initiate this process, we identified promoter peaks as scATAC peaks located within ±1,000 bp upstream and downstream of a TSS. Peaks co-accessible to these promoter peaks were then identified using the *getCoAccessibility()* function, applying a *correlation cutoff of 0.5* and a *resolution of 1*, which were default parameters.

Predicted target genes for each snATAC peak were generated using the *getPeak2GeneLinks* function. This step integrated barcode-matched RNA expression data from snRNA-seq, incorporating a correlation cutoff of 0.45, an *FDR cutoff of 1e-04*, and a *resolution of 1*. By integrating co-accessibility with gene expression information, this analysis enables the identification of potential regulatory interactions and enhancer-gene associations, thereby providing insights into the functional implications of chromatin accessibility in driving gene expression patterns. We have developed a Shiny App where the browser tracks can be visualized (https://tnbcworkbench.org/shiny/ShinyArchRUiO).

### Trajectory analysis

To unravel the molecular processes initiated by transplantation that drive changes in cell states, we embarked on pseudo-time or trajectory analysis spanning all clusters. The root cluster, representing adult basal cells (the transplanted cell type), was selected to initiate this exploration. In our pursuit of unbiased lineage trajectories, we opted for Monocle trajectory analysis, given that ArchR necessitates a user-defined trajectory backbone to establish a preliminary ordering of cell groups or clusters.

To construct an encompassing trajectory, we employed the *getMonocleTrajectories()* function. This function’s principal group was set as the adult basal cluster (Ba.adult), and specific parameters, including use_partitions and close_loop (both set to false), were specified. By not calculating closed loops and learning a single graph across all partitions, we ensured the generation of an overall trajectory that best represented the underlying cell transitions.

Subsequently, we pinpointed 10 pair-wise transitions from this global trajectory across diverse cell types. These transitions allowed us to delineate pseudotime trajectories for each pair individually. We also generated a 3D plot for the overall trajectory to help us visualize the clusters that were overlapping. This trajectory analysis utilizing Monocle provides valuable insights into the orchestrated progression of cell states post-transplantation, contributing to our understanding of the underlying molecular dynamics and potential programs that are set in motion by the transplantation process.

### Integrative pseudo-time analyses

To dissect the drivers behind cellular transitions, we executed integrative analyses, notably involving the identification of positive TF regulators through correlation between gene expression and motif accessibility changes across pseudo-time for all 12 transitions (T1-T12). These transition-driving TFs exhibited a positive correlation between their gene expression and the changes in accessibility of their corresponding motifs across the diverse transitions. ArchR facilitated this identification process by stratifying motifs based on their variation across clusters using chromVAR deviation-z scores.

To conduct this analysis, we employed the *correlateTrajectories()* function with default parameters (*corCutOff = 0.5*). This function ingests trajectory objects obtained from the *getTrajectories()* function, which encapsulate transitioning cell expression values from the GeneExpressionMatrix, along with deviation scores from the MotifMatrix. A critical step is the computation of a "variance quantile”, a normalized metric for a given feature, enabling the derivation of correlations across disparate assays.

To obtain significant correlating TFs, we employed a combined *"Variance Quantile Cutoff" > 0*. The identified TFs were visualized using a heatmap generated with the *plotTrajectoryHeatmap()* function. This allowed us to unravel the key transcriptional regulators driving cellular transitions across multiple transitions, providing valuable insights into the orchestrated changes in cell states post-transplantation.

### Differential expression gene and functional enrichment analyses

Differential Expression Analysis (DEA) was done for every pair of clusters that constitute the source and destination of a transition (e.g. for T1, the source Ba.adult and destination D3-5.CyclingBa+D3-5.Ba/LS) with cutoffs (*FDR ≤ 0.05* and *abs(Log2FC) ≥ 0*). On each of the DEGs, we performed gene set enrichment and pathway analysis using hyper-geometric enrichment. Up and down regulated genes were identified based on cutoffs (*log2FC > 0* and *log2FC < 0*, respectively).

Pathway analyses were performed using Hallmark with msigdbR, Mouse MSigDB Collections^70^, Reactome with ReactomePA^71^, and GO Biological Processes databases with the clusterProfiler^72^. The results were visualized using *ggplot2* and *clusterProfiler* packages.

### Mammary organoids for drug screening

Sorted basal cells (1000 cells) were encapsulated in Matrigel domes (15 μl, Corning, 356231). After gelification at 37°C was achieved, complete media (350 μl) was added on top. Complete media was adapted from previously reported^73^, and consisted of Advanced DMEM/F-12 (Gibco, 12634-010), Glutamax (1x, Gibco, 35050-061), Hepes (10 mM, Gibco, 25534), Pen/Strep (1%, Gibco, 15140-122), R-spondin (500 ng/ml, R&D Systems, 4645-RS), Noggin (100 ng/ml, Peprotech, 250-38-50), B27-Supplement (1x, Gibco, A35828-01), N-acetylcysteine (1.25 mM, Sigma, A9165), Nicotinamide (10 mM, Sigma, NO636), Y27632 (5 μM, AdipoGen AG-CR1-3564), A83-01 (500 nM, Tocris, 2939), SB202190 (1 μM, Cayman Chemical, 10010399), FGF-7 (5 ng/ml, Peprotech, 100-19), FGF-10 (20 ng/ml, Peprotech, 100-26), Neuregulin (5 nM, Peprotech, 100-03), and mEGF (5 ng/ml, Thermo Fisher Scientific, PMG8041). The following inhibitors were added to the media: Roscovitine (or Seliciclib, 20 μM, MedChemExpress, HY-30237), Palbociclib (2 μM, MedChemExpress, HY-50767), Nutlin-3a (10 μM, MedChemExpress, HY-10029), Nutlin-3b (10 μM, MedChemExpress, HY-15335), Cyclopamine (25 μM, MedChemExpress, HY-17024), Sonidegib (5 μM, MedChemExpress, HY-16582A), UVI 3003 (50 μM, MedChemExpress, HY-107500), SR11302 (50 μM, MedChemExpress, HY-15870), JSH-23 (50 μM, MedChemExpress, HY-13982), Adalimumab (5 μg/ml, MedChemExpress, HY-P9908), QNZ (or EVP4593, 5 μM, Selleck Chemical, S4902), Galunisertib (50 μM, Selleck Chemical, S2230), Avagacestat (10 μM, R&D Systems, 6363), LDN193189 (2 μM, MedChemExpress, HY-12071), or DMSO to serve as vehicle control. Media was changed after 4 days with replenishment of the drugs.

### Quantification and statistical analysis

All statistical analyses were performed using GraphPad Prism 9.5.0 (GraphPad Software, La Jolla, CA). All values are expressed as mean +/- standard error. Statistical significance was calculated by either unpaired Student’s t-tests assuming two-tailed distribution and unequal variances, one-way ANOVA with Tukey’s multiple comparison test, or two-way ANOVA with Sidak’s multiple comparison test. The value of n refers to the number of mice used in a given study. A p value of ≤ 0.05 was considered statistically significant. For all figures, asterisks denotes statistical significance and p values are denoted in the figure legends.

